# Pre-determined diversity in resistant fates emerges from homogenous cells after anti-cancer drug treatment

**DOI:** 10.1101/2021.12.08.471833

**Authors:** Yogesh Goyal, Ian P. Dardani, Gianna T. Busch, Benjamin Emert, Dylan Fingerman, Amanpreet Kaur, Naveen Jain, Ian A. Mellis, Jingxin Li, Karun Kiani, Mitchell E. Fane, Ashani T. Weeraratna, Meenhard Herlyn, Arjun Raj

## Abstract

Even amongst genetically identical cancer cells, therapy resistance often only emerges from a very small subset of those cells. Much effort has gone into uncovering the molecular differences in rare individual cells in the initial population that may allow certain cells to become therapy resistant; however, comparatively little is known about variability in the resistant outcomes themselves. Here, we develop and apply FateMap, a framework that combines DNA barcoding with single-cell RNA sequencing to reveal the fates of hundreds of thousands of clones exposed to anti-cancer therapies. We show that resistant clones emerging from single-cell-derived cancer cells adopt molecularly, morphologically, and functionally distinct fate types. These different resistant types are largely predetermined by molecular differences between cells before addition of drug and not by extrinsic cell-specific microenvironmental factors. Changes in dose and kind of drug can, however, switch the resistant fate type of an initial cell, even resulting in the generation and elimination of certain fate types. Diversity in resistant fates was observed across several single-cell-derived cancer cell lines and types treated with a variety of drugs. Cell fate diversity as a result of variability in intrinsic cell states may be a generic feature of response to external cues.

## Introduction

Individual cells within a population can respond to signals and stresses in different ways. Such differences often arise from intrinsic differences in the cells present before the signal or stress is applied, and a growing body of evidence suggests that these differences can be non-genetic in origin (Chen and Larson, 2016; Raj and van Oudenaarden, 2008; Symmons and Raj, 2016). Hence, considerable effort over the past twenty years has been put into characterizing the variability of the molecular profiles of individual cells in a population (Elowitz et al., 2002; Gupta et al., 2011; Kinker et al., 2020; Raj et al., 2006; Rodriguez et al., 2019; Shakiba et al., 2019; Sharma et al., 2010; Spencer et al., 2009; Wouters et al., 2020). More recently, the development of high-throughput strategies for single-cell barcoding have enabled us to track differences in initial molecular states through to different outcomes after the application of signals and stresses (Bhang et al., 2015; Biddy et al., 2018; Emert et al., 2021; Frieda et al., 2017; Gutierrez et al., 2021; Leighton et al., 2021; Oren et al., 2021; Rodriguez-Fraticelli et al., 2020; Tian et al., 2021; Umkehrer et al., 2021; Weinreb et al., 2018). However, far less attention has been paid to characterizing variability in the outcomes themselves. Typically, the implicit assumption is that these outcomes are binary: induced or not induced, proliferative or nonproliferative, alive or dead. It is possible, however, that there is a far richer set of outcome states than presumed.

An important example of variable response to external stress is the development of therapy resistance in cancer. While many drugs are successful at killing the vast majority of cancer cells, they often leave behind a small subpopulation that is drug-resistant and resumes proliferation, which represents a major barrier to cures. Recent work from our group and others has revealed that even single-cell-derived populations of cancer cells harbor these subpopulations, which are marked by fluctuations in the expression of particular genes, including several receptor tyrosine kinases (Marin-Bejar et al., 2021; Pillai and Jolly, 2021; Rambow et al., 2018; Roesch et al., 2010, 2013; Schuh et al., 2020; Shaffer et al., 2017, 2020; Su et al., 2017; Wouters et al., 2020). When drug is applied to these single-cell-derived populations, these drug-resistant cells grow out into resistant colonies, forming a resistant population composed of a relatively small number of resistant clones. Are all the cells in this resistant population the same, both in terms of their molecular profile and behavior? If individual resistant cells are different from each other, to what extent does the clonal structure of the population explain this variability—i.e., does each clone harbor all resistant cell types, or are the cells within each clone relatively homogeneous and is most of the variability across clones? While different resistance mechanisms have been documented in the literature (Krepler et al., 2017; Marin-Bejar et al., 2021; Rambow et al., 2018; Ramirez et al., 2016; Su et al., 2017; Tirosh et al., 2016), it is unclear whether different resistant cell types can emerge from a homogeneous initial population. Differences in proliferative capacity of clones suggest some degree of heterogeneity between resistant clones (Emert et al., 2021; Oren et al., 2021; Umkehrer et al., 2021), but little is known about the heterogeneity of resistant populations and its origins, both at the molecular and phenomenological level.

To address this gap, we developed FateMap, a framework combining single-cell RNA sequencing, DNA barcoding, and computational analysis to follow the fates of thousands of individual cancer cell clones as they acquire resistance. Our work reveals that when treated with targeted therapies, single-cell-derived cancer cells—expanded from a single cell and grown in homogeneous conditions—can give rise to molecularly and functionally diverse resistant fates. The specific resistant fate that cells adopt is strongly predetermined by the intrinsic differences between them preceding drug exposure, and is independent of extrinsic (e.g. neighboring cells) factors. Varying the dose or type of drug can change the mapping between intrinsic differences and resistant fate, resulting in characteristic fate switching and an altered number and proportion of distinct fates. For example, treatment with trametinib, a MEK inhibitor, eliminated a specific resistant fate that vemurafenib, a BRAF inhibitor, could not. Transcriptional and functional diversification and the predetermination of distinct resistance fates is consistent across cancers (melanoma and triple-negative breast cancer) and therapies (targeted therapy and cytotoxic chemotherapy). Altogether, our results demonstrate that a rich set of outcomes can arise in response to external stresses due to pre-existing variability within a seemingly homogeneous initial population of cells.

## Results

### Diverse transcriptional resistant fates emerge following targeted therapy treatment of single-cell-derived cancer cells

Molecular differences between genetically identical cancer cells can result in some cells surviving treatment with targeted therapies and developing resistance. We wondered whether the resistant population of cells that emerged from the treatment of single-cell-derived cancer cells all adopted similar or distinct fates. To test this idea, we focused on BRAF^V600E^-mutated melanoma, where treatment of single-cell-derived cells with targeted therapy drug vemurafenib leads to survival of a rare (1:1,000 or even more rare) subpopulation of cells, which then begin to proliferate to form resistant colonies (also referred to as clones) (Figure 1A; Supplementary Movie 1). To test whether the resistant fates of cells were similar, we performed single-cell RNA sequencing on a mixture of all the resistant colonies that emerged on a tissue culture dish. We found that the resistant fates exhibited extensive diversity in their gene expression profiles (Figure 1B; Supplementary Figure 1A,B,E). As expected, we found subpopulations of cells that expressed canonical resistance markers (e.g. *AXL, SERPINE1*), but we also found several other subpopulations that expressed their own distinct sets of marker genes. Often, these subpopulations would express multiple markers reminiscent of particular cell types; for example, smooth muscle (e.g. *ACTA2, ACTG2, MYOCD*), neural crest (e.g. *NGFR, S100B, GAS7*), adhesive (e.g. *VCAM1, PKDCC, ITGA8*), melanocytic (e.g. *MLANA, SOX10, MITF*), or type-1 interferon signaling-enriched (e.g. *IFIT2, DDX58, OASL*) (Figure 1C; Supplementary Figure 1A,B). Thus, it was clear that diverse resistant cell states can emerge from single-cell-derived cancer cells upon treatment with targeted therapy.

**Figure 1:**
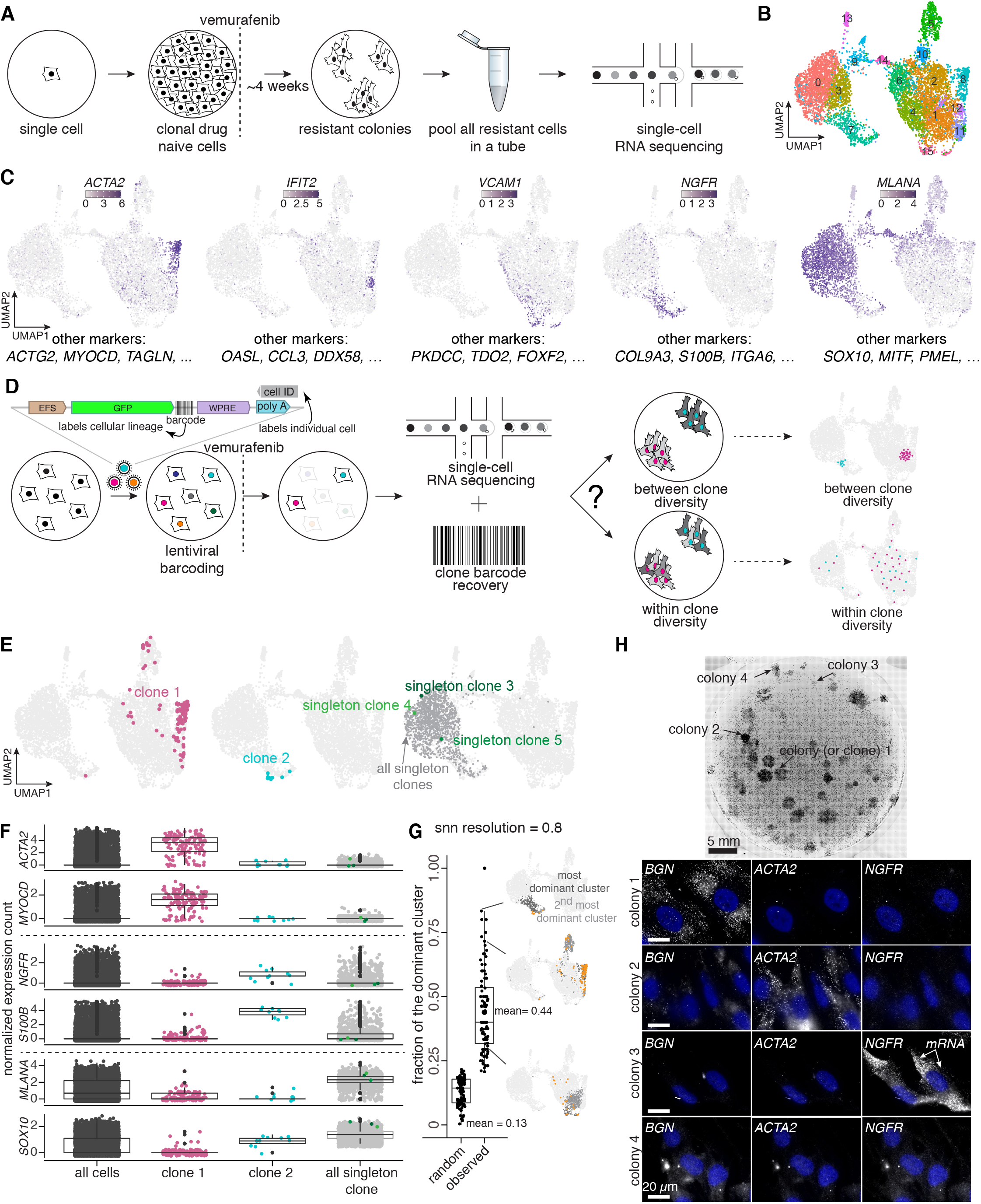
FateMap reveals that between-clone fate diversity arises from a single cell upon therapy treatment. a. Schematic of the experimental design where we exposed single-cell-derived WM989 A6-G3 melanoma cells to the targeted therapy drug vemurafenib, resulting in resistant colonies in 3-4 weeks. We then mixed the resistant colonies together and immediately performed single-cell sequencing on the resistant population. b. We applied the Uniform Manifold Approximation and Projection (UMAP) algorithm within Seurat to the first 50 principal components to visualize differences in gene expression. Cells are colored by clusters determined using Seurat’s FindClusters command at a resolution of 0.6 (i.e. “Seurat clusters, resolution = 0.6”). c. On the UMAP, we recolored each cell by its expression for a select subset of genes that were identified as differentially expressed via the Seurat pipeline and marked different clusters. Additional gene names with similar UMAP expression profiles are provided below individual UMAP panels. *ACTA2*, which marks smooth muscles, is found largely in Seurat cluster 8; *IFIT2*, which marks type-1 interferon signaling, is found largely in cluster 12; *VCAM1*, which marks vascular adhesion, is found largely in cluster 15; *NGFR*, which marks neural crest cells, is found largely in cluster 7; *MLANA*, which marks melanocytes, is found largely in clusters 0 and 3. d. Schematic of FateMap for labeling WM989 A6-G3 melanoma cells with unique DNA barcodes prior to exposure with vemurafenib. For the experiment shown, we transduced WM989 A6-G3 cells at an MOI of ~0.15 with a barcode library similar to that described in (Emert et al., 2021). We exposed the barcoded cells to vemurafenib for 3-4 weeks and performed single-cell RNA sequencing and barcode sequencing on the resultant colonies. We asked whether the resistant cells sharing a barcode (a resistant clone) were more transcriptionally similar to each other than other clones. e. Three examples to demonstrate that a clone (cells sharing the same barcode) is constrained largely in a specific transcriptional cluster such that cells within a clone are more transcriptionally similar to each other than cells in other clones. Some clones are larger in size than others, and some exist as singletons, meaning they survive vemurafenib treatment but do not necessarily divide in the drug. f. We compared a selected subset of two genes corresponding to each example covered in e. We found that clones from distinct regions/clusters of the UMAP are enriched for expression of specific genes that characterize those regions. g. We quantified the preference for a specific cluster across all barcode clones (clone size>4). Specifically, we calculated the fraction of dominant clusters for each clone and found it to be significantly higher than that for randomly selected cells. While the analysis plotted here is for a cluster resolution of 0.8, we show that this result is consistent across a range of resolutions (0.4 to 1.2; Supplementary Figure 2). Representative examples of clones covering different fractions of dominant clusters. h. We performed RNA FISH for a subset of genes that marked different single-cell resistant populations. Consistent with the results from FateMap, we found resistant colonies that were selectively positive for each of the three markers tested (*NGFR, ACTA2, BGN*), and ones that were negative for all of them.

After adding therapy, resistant cells grow out as separate clones from individual rare cells amidst the initial drug-naive population (Supplementary Movie 1). Thus, the question arises as to whether the clones perfectly match the different transcriptional subtypes revealed by the single-cell RNA sequencing. Specifically, we wondered whether each clone harbored several if not all of the transcriptionally diverse cell types or if each clone was characterized by its own distinct transcriptional profile (Figure 1D). Within-clone and between-clone transcriptional diversity each imply very different sources of the observed transcriptional variability. The former would imply that resistant cells can dynamically switch between these cellular states *after* a particular clone has become resistant. The latter would suggest that resistant clones are inherently transcriptionally stable, at least in terms of the states observed by single-cell RNA sequencing. In order to differentiate between these scenarios, we needed a way to simultaneously transcriptionally profile individual resistant cells while determining which resistant clone each cell came from.

To this end, we developed FateMap, a method that uses transcribable DNA barcoding to identify clones from thousands of resistant cells in a single experiment (Figure 1D). Briefly, FateMap begins with lentiviral integration of barcodes into the DNA of therapy-naive cells. With a large barcode library complexity (~59 million unique barcodes, see Methods; and Supplementary Figure 1C) and small multiplicity of infection (MOI), we ensured that many thousands of cells could be uniquely barcoded, which was critical for following the rare cells that become resistant. The DNA barcodes lie in the 3’ untranslated region of GFP, and the entire gene including the barcode is integrated into the cells’ DNA. After barcoding, we sorted the GFP-positive cells to enrich for barcoded cells, exposed them to vemurafenib, collected the resistant populations, performed single-cell RNA sequencing, and extracted the FateMap barcode (clone information) for each cell. Our approach relied on the fact that the single-cell RNA sequencing cDNA library contained the GFP mRNA with both the clone barcode information and the cell identifier, which we selectively amplified and sequenced to connect the clone and the single-cell transcriptome information (see Methods). In this way, FateMap enabled simultaneous extraction of transcriptomic profile and clone identity information for individual cells within the resistant population.

Armed with these data, we could identify which cells on the UMAP originated from the same initial resistant cell. We found that clones fell predominantly within a constrained region of the UMAP space marked by distinct transcriptional markers (Figure 1E). Therefore, the transcriptionally variability we observed across the entire resistant population was dominated by variability across resistant clones, and not within individual clones. For example, a large and a small resistant clone was enriched for genes expressed in smooth muscle (e.g. *ACTA2, MYOCD*) and neural crest cells (e.g. *NGFR, S100B*), respectively (Figure 1E,F). There were also proliferating clones that were enriched for canonical resistance markers (e.g. *AXL, SERPINE1*) (Supplementary Figure 1B,D). We also found a subpopulation of cells that survived therapy but appeared to be largely non-proliferative; i.e. these clones consisted of only one barcoded cell detected (which we refer to as singleton) and were enriched for genes expressed in melanocytes (e.g. *SOX10, MLANA*) (Figure 1E,F) (98.6% of all clones within clusters 0 and 3 were singletons). It is worth noting that these cells might normally be missed in conventional analyses of therapy-resistance due to their rarity. (In some cases, the cells in a clone would come from two non-neighboring clusters, e.g., cluster 15 and 6 for three clones marked by genes *VCAM1* and *APOE*, respectively (Supplementary Figure 1F,G). These constituted a minority of cases.)

To more formally quantify the degree to which each clone was transcriptionally homogeneous, we calculated, for each clone, the fraction of cells from the most dominant transcriptional cluster (clusters are identified from the neighbor graph on the principal component space) within the clone (e.g., 72% of cells from clone 1 belonged to cluster 8 at a cluster resolution of 0.6) (Figure 1E,G). As a baseline for comparison, we randomly sampled groups of cells matching the size of each observed clone (“pseudo-clones”) from the single-cell RNA sequencing dataset and similarly calculated the fraction of the most dominant cluster. We found that the mean fraction of the dominant cluster for the experimentally observed clones was 3.4-fold-higher than the randomly sampled clones (mean experimentally observed fraction = 0.44 and mean randomly sampled fraction = 0.13) (Figure 1G; Supplementary Figure 2A-B), showing that clones largely consisted of transcriptionally similar cells. This conclusion held for a large range of resolution values to identify clusters using the shared nearest neighbor (SNN) modularity optimization based clustering algorithm within Seurat (Supplementary Figure 2C-E).

We wanted to confirm whether our sequencing based FateMap approach correctly captured and matched distinct transcriptional fates and clones with an independent method. We designed single molecule RNA FISH probes for a subset of genes (*ACTA2, NGFR, BGN*) that were differentially expressed and belonged to distinct clusters (Figure 1C; Supplementary Figure 1E), and performed multiplexed RNA imaging on a large plate containing several resistant clones. We verified that the selected markers indeed were expressed only in distinct resistant clones, and that for some clones none of the three markers were expressed (Figure 1H), thus validating the results we obtained by FateMap.

### Gene expression differences between clones correspond to differences in morphology, proliferation, and invasiveness

We wondered whether transcriptionally distinct resistant clones had distinct phenotypic properties. We first focused on differences in proliferative capacity between resistant clones, given that resistant colonies are often differently sized upon visual inspection (Emert et al., 2021) (Figure 1E,H). We obtained the relationship between transcriptional profiles and proliferative abilities of resistant clones from FateMap by measuring the cell count per clone for distinct transcriptional clusters (Figure 2A,B; Supplementary Figure 1H). We found that distinct clusters had different proliferative capacities. Some clusters, e.g. those marked by *ACTA2, AXL*, and *VCAM1*, tended to primarily form large colonies (Figure 2B), while those marked by *NGFR* (cluster 7) tended to form small colonies and singletons (Figure 2B). Similarly, the *MLANA*-high resistant fate clusters (clusters 0 and 3) were primarily composed of singletons, suggesting minimal proliferative capacity in these cells (Figure 2B).

**Figure 2:**
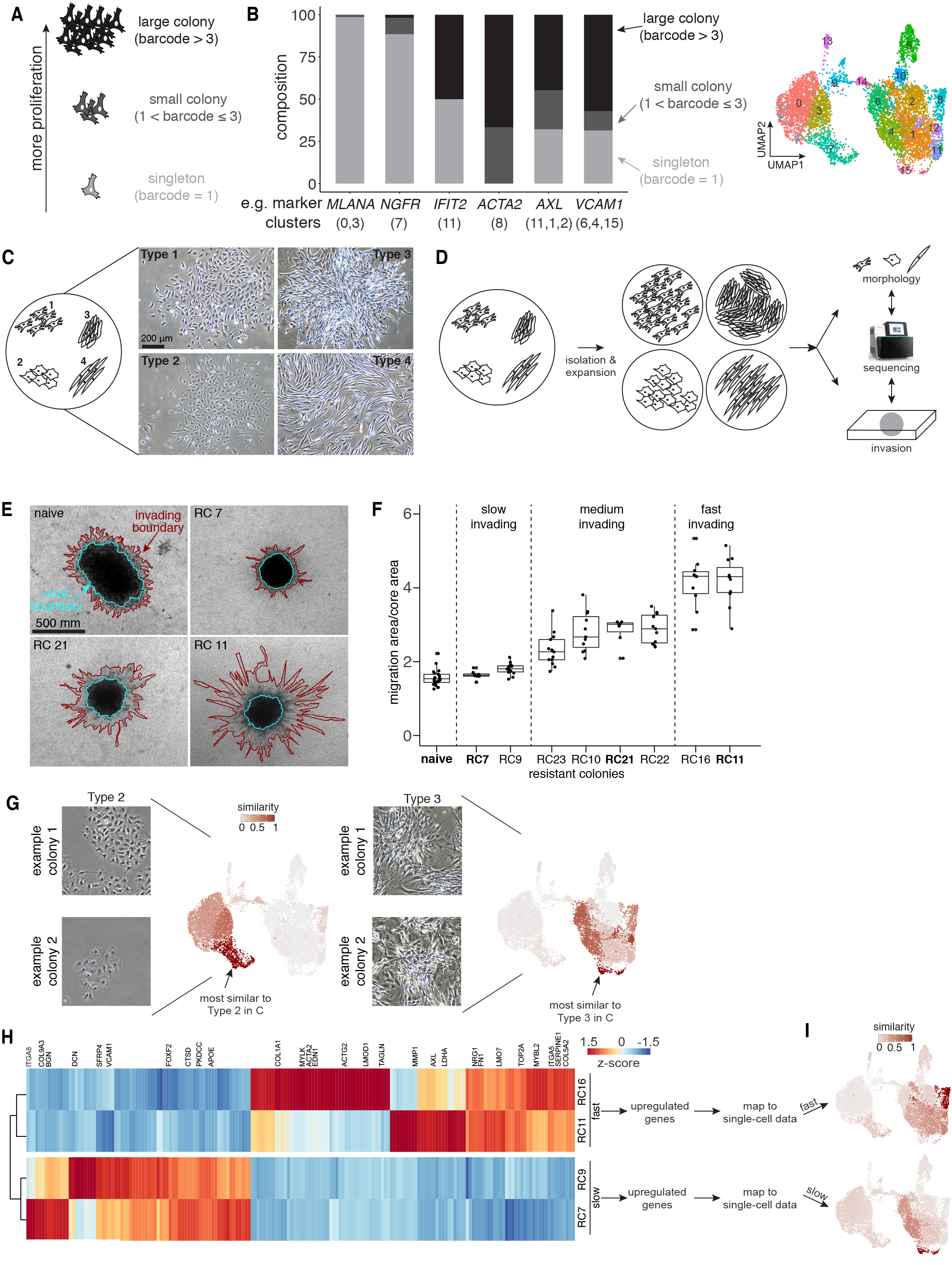
Gene expression differences between clones correspond to differences in morphology, proliferation, and invasiveness. a. We classified colonies into three groups: those that only exist as singletons (i.e. they survive but do not proliferate), small colonies, and large colonies. b. (left) Markers for different clusters exhibited different proliferative capacities. For example, clusters 0 and 3, which are enriched for the gene *MLANA*, predominantly (98.6%) contained singletons. Cluster 7, enriched for the gene *NGFR*, was largely composed of singletons or small colonies. Other markers, including ACTA2, AXL, and VCAM1, which belonged to different clusters, primarily contained small and large colonies, and relatively fewer singletons. (right) UMAP where cells are colored by clusters determined using Seurat’s FindClusters command at a resolution of 0.6 (i.e. “Seurat clusters, resolution = 0.6”). c. (left) Traced representative cells in Adobe Illustrator and created cartoon schematics based on visual inspection of orientation and density. (right) Brightfield images of resistant colonies exhibiting different types of morphologies. d. Schematic of isolation and expansion of resistant colonies emerging from either vemurafenib or trametinib treatment belonging to different morphology types, (a subset of) which were then bulk RNA sequenced, categorized for morphology, and measured for invasiveness. e. A spheroid assay in which cells from resistant colonies were seeded at 3000 cells/well and allowed to form spheroids over 96 to 120 hours. Those that formed 3D aggregates (spheroids) were then embedded in a collagen matrix. Red marks the invading boundary and blue marks the core of the embedded spheroid. f. Quantification of invasiveness of resistant colonies emerging from treatment with trametinib. We measured and ranked the spheroid invasiveness by computing the ratio of the area under the invading boundary (red) and the embedded spheroid core (blue). Each dot represents one spheroid with each condition having multiple spheroids. g. Mapping of morphology onto the single-cell RNA sequencing dataset from FateMap by comparing genes differentially expressed from each morphology compared to all other resistant colonies, with the UMAP colored for the similarity score. Similarity score represents the degree of overlap of differentially expressed genes between bulk-sequencing data and each cluster. Resistant colony type 2 maps predominantly to cluster 7 (and to some extent 3), while type 3 maps to cluster 15 (and to some extent 4,6, and 12). h. Mapping of invasiveness onto the single-cell RNA sequencing dataset from FateMap by comparing genes differentially expressed between the two slowest and the two fastest invading resistant colonies, with the UMAP colored for the similarity score. The slowest invading colonies have a high similarity score for cluster 15 (and to some extent 4 and 6), while the fastest invading colonies have a high similarity score for cluster 8 (and to some extent 1).

Next, we asked whether different resistant clones exhibited distinct morphologies. We performed brightfield imaging of resistant colonies on the plate and identified several distinct morphologies (Figure 2C), including colonies with cells that appeared epithelial (type 1), cells that grew slower and were more transparent (type 2), cells that tended to grow primarily on top of each other (type 3), and elongated cells (type 4). For a subset of the types, we were able to isolate the colony and perform multiple cycles of growth and replating; for these types, the colonies retained their morphology (Figure 2D) (some colonies did not survive the replating process).

Given the morphological distinctions between the resistant colonies, suggesting differences in their cell biological properties, we then tested whether they have differences in invasive potential. We manually isolated 64 of therapy resistant colonies from multiple parallel experiments and expanded them for months (some colonies did not survive the expansion process). We then used a spheroid assay which measures the invasiveness of cancer cells in the collagen matrix (Kaur et al., 2019). We formed 3D aggregates (spheroids) from a subset of resistant colonies (with sufficient cell numbers), embedded them in a collagen matrix, and measured their invasiveness by looking for the area under invading boundary (red) relative to that of the embedded spheroid core (blue) (Figure 2E,F). We found that different resistant clones emerging from the same condition resulted in dramatically different invasion areas in the collagen matrix (some colonies were unable to aggregate into spheroids) (Figure 2E,F). Therefore, morphologically and functionally distinct resistant clones can emerge from a single cell upon treatment with therapy drugs.

Given the observed variation in morphology and invasiveness, we wonder whether we could connect these phenotypic differences to distinct transcriptional profiles (Figure 2D). We performed bulk RNA sequencing of manually isolated colonies with known morphology and invasive potential. We identified genes differentially expressed between morphology types from bulk RNA sequencing and used these gene sets to map morphologies to single-cell clusters from Fatemap (see Methods) (Figure 2G). For example, we found the type 2 and type 3 morphology from Figure 2C corresponded to gene expression signatures most similar to the *NGFR*-high cluster (cluster 7) and *VCAM1*-high cluster (cluster 15) (Figure 2G; Supplementary Table 4). Similarly, we found several genes differentially expressed between slow and fast invading resistant colonies from bulk sequencing which we were able to connect with the UMAP clusters from FateMap (Figure 2H). For example, we found that the fastest invading resistant colonies were enriched for expression of genes that mark cluster 8, including *ACTA2, TAGLN* and *EDN1* (Figure 2H). Therefore, gene expression differences between clones correspond to functional differences in proliferation, morphology, and invasiveness.

### Diverse drug fates are observed in resistant clones from another melanoma cell line and from breast cancer cells

We asked whether the diversity of resistant clones was specific to the melanoma cell line we studied (WM989 A6-G3) or if it was a more general property of resistant cancer cells. We first checked whether other melanoma cell lines showed similar diversity in resistant cell states by performing FateMap on a single-cell-derived clone of another patient-derived melanoma cell line (WM983B E9-C6) that we also treated with vemurafenib. Brightfield imaging of the resistant clones emerging from vemurafenib treatment revealed morphological and proliferative differences between resistant clones, similar to what we observed for WM989 A6-G3 cells (Supplementary Figure 3A). Applying FateMap to these resistant populations of cells, we similarly observed single-cell transcriptional diversity (Supplementary Figure 3B,C). Moreover, we observed that many transcriptional signatures overlapped with those observed in WM989 A6-G3 cells, including neural crest-like markers (e.g. *NGFR, S100B*), melanocytic markers (e.g. *MLANA, SOX10*), type-1 interferon signaling markers (e.g. *IFIT2, DDX58*), and general resistance-associated markers (e.g. *AXL, VCAM1*) (Supplementary Figure 3C). Analysis of individual resistant clones via barcoding also revealed a similar diversity in their proliferative properties as observed in the WM989 A6-G3 line. We again found that resistant clones were present in many different sizes, from singletons to small colonies to large colonies, and the clones typically demarcated particular regions of transcriptional space (Supplementary Figure 3D, E). Both the neural crest-like clones (*NGFR*-high) and the melanocytic-like clones (*MLANA*-high) appeared to be either singletons or small colonies, also similar to what we observed for WM989 A6-G3 cells (Supplementary Figure 3D-F).

Next, we wondered if other cancers subjected to therapies with different mechanisms of action would also give rise to a diversity of resistant cell fates. We used another single-cell-derived cell line, MDA-MB-231-D4 (Shaffer et al., 2020), which is a triple negative breast cancer cell line (i.e., does not express high levels of *HER2*, estrogen receptors, or progesterone receptors). Treatment of these cells with the chemotherapy drug paclitaxel, a common line of treatment for such breast cancers, killed most of the cells, leaving behind rare cells (~1:1000, see Methods) that proliferated to form resistant clones (Supplementary Figure 4A). Similar to the melanoma cell lines, brightfield imaging revealed that the morphologies of cells were different between drug-resistant colonies, and that the size of the resistant colonies varied widely (Supplementary Figure 4B,C). Applying FateMap to the resistant population of MDA-MB-231-D4 cells revealed transcriptional diversity in resistant fates (Supplementary Figure 4D). Analyzing clones using barcodes, we found that certain clones were far more proliferative than others (Supplementary Figure 4F,G). The cells comprising clones were not as clearly confined to particular transcriptional clusters as in the melanoma cell lines (Figure 1E), but they were still more confined than random (Supplementary Figure 4F-H), and different clones still had distinct molecular expression profiles (Supplementary Figure 4F-H). This smaller degree of within-clone transcriptional homogeneity may be a property of the cell type (breast cancer vs melanoma) and/or type of treatment (chemotherapy vs. targeted therapy). Together, our results establish that the emergence of cell fate diversity upon treatment of single-cell-derived cancer cell lines may be universal.

### Mouse xenograft models confirm transcriptional diversity and its spatial patterns

The more complex microenvironments present *in vivo* may alter the transcriptional diversity we observed in the tissue culture dish. To test whether that was the case, we used cells from another single-cell-derived clone of WM989 cells, WM989 A6-G3 5a3 (first used in (Torre et al., 2021)), injected them subcutaneously into mice, subjected the mouse to targeted therapy treatment PLX4720, harvested the resistant tumors, and measured single cell gene expression of markers via RNA FISH (Figure 3A). Upon performing large-scale scans of tumor tissue sections, we found that gene markers for the various resistant fates we observed in cell culture (e.g. *AXL, NGFR, SOX10, BGN, ACTA2*) were present in distinct regions of the tumor sections (Figure 3B,C). In contrast, the expression patterns of *UBC*, a housekeeping gene, remained relatively homogeneous (Figure 3C), similar to what we observed from FateMap on cells in a tissue culture dish (Supplementary Figure 1I). Thus, the diversity of resistant fates observed in cell culture can also appear in more realistic microenvironments, raising the possibility that it may be an inherent property of the cells.

**Figure 3:**
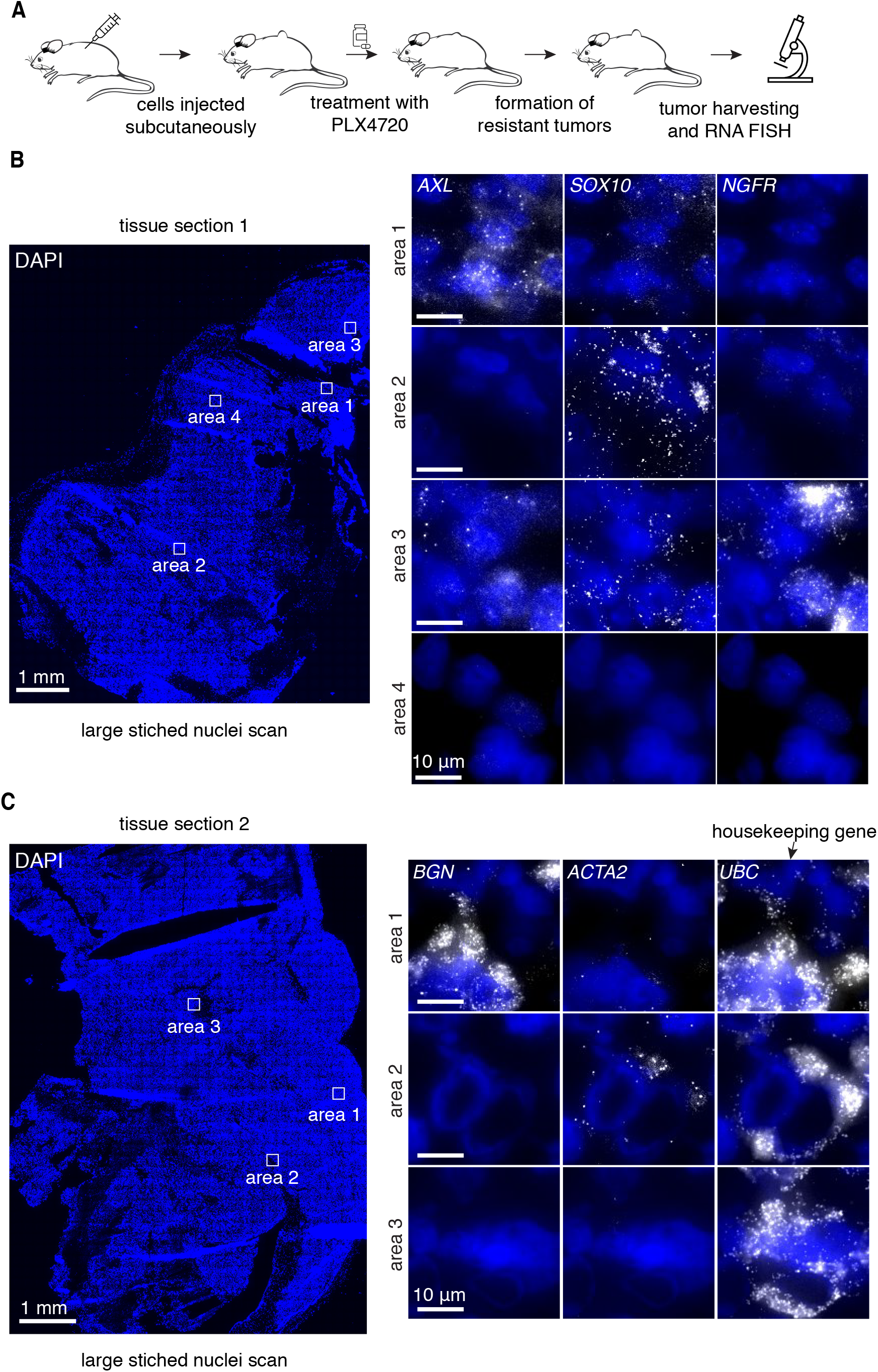
Mouse xenograft models confirm transcriptional diversity and its spatial patterns. a. We used cells from another single-cell-derived cell line from WM989 A6-G3 cells, WM989 A6-G3 5a3 (Torre et al., 2021), injected them subcutaneously into mice, subjected the mouse to targeted therapy, harvested the resistant tumors and sectioned them, and performed large RNA FISH tissue scans. b. (left) A large-scale scan for DAPI-stained nuclei of a resistant tumor tissue section. (right) RNA FISH imaging of three gene markers (*AXL, NGFR, SOX10*) from four different areas on the tissue section. Each of these markers were selectively expressed in distinct regions of the sections, with some regions lacking all three markers. c. (left) A large-scale scan for DAPI-stained nuclei of another resistant tumor tissue section. (right) RNA FISH imaging of three gene markers (*BGN, ACTA2, UBC*) from three different areas on the tissue section. While *BGN* and *ACTA2*, two differentially expressed genes observed in FateMap, were selectively expressed in distinct regions of the sections, the housekeeping gene *UBC* was relatively homogeneous.

### Cells are predestined for distinct resistant fates upon exposure to targeted therapy and chemotherapy

We wondered whether the transcriptional and phenotypic variability in therapy resistant clones was the result of intrinsic differences in the molecular expression states of cells preceding drug exposure. Alternatively, the resistant fate may be determined extrinsically, e.g. by the location and immediate neighbors of cells (Jiang et al., 2021). An “identical twin” analysis combined with Fatemap enabled us to distinguish between these possibilities.

Briefly, upon uniquely barcoding cells, we allowed them to divide for a few divisions and then separated the population into two equal split populations A and B (“twins”) such that most barcoded clones (>90%) were present in each group (Figure 4A). We then applied the vemurafenib to both split populations and determined their resistant fates that emerged from each split population. We reasoned that if the resistant fate of a cell was intrinsically predetermined, then its twin would share the same fate (assuming that the predetermination has enough memory to be maintained over at least a few cell divisions (Emert et al., 2021; Schuh et al., 2020; Shaffer et al., 2017, 2020). We first asked whether the barcodes that showed up in the resistant population from split population A also showed up in the resistant population from split population B, which would be consistent with intrinsic predetermination of outcome. Similar to our approach in a recent study (Emert et al., 2021), we extracted the genomic DNA (gDNA) from each split population, amplified and sequenced the DNA barcode region. Additionally, prior to the library preparation, we added specific amounts of known barcoded cells as standards (see Methods), which enabled us to convert sequencing reads to the cell numbers, thus enabling comparisons of absolute number of cells among clones both within and across the split populations A and B. We found that the ability to survive and proliferate (including the degree of proliferation) was strongly correlated between barcodes across the two groups and significantly more likely than random (Figure 4B,C). Thus, the property of being resistant and the phenotypic degree of resistance appeared to be encoded in the intrinsic state of the pre-resistant cell before therapy was even applied.

**Figure 4:**
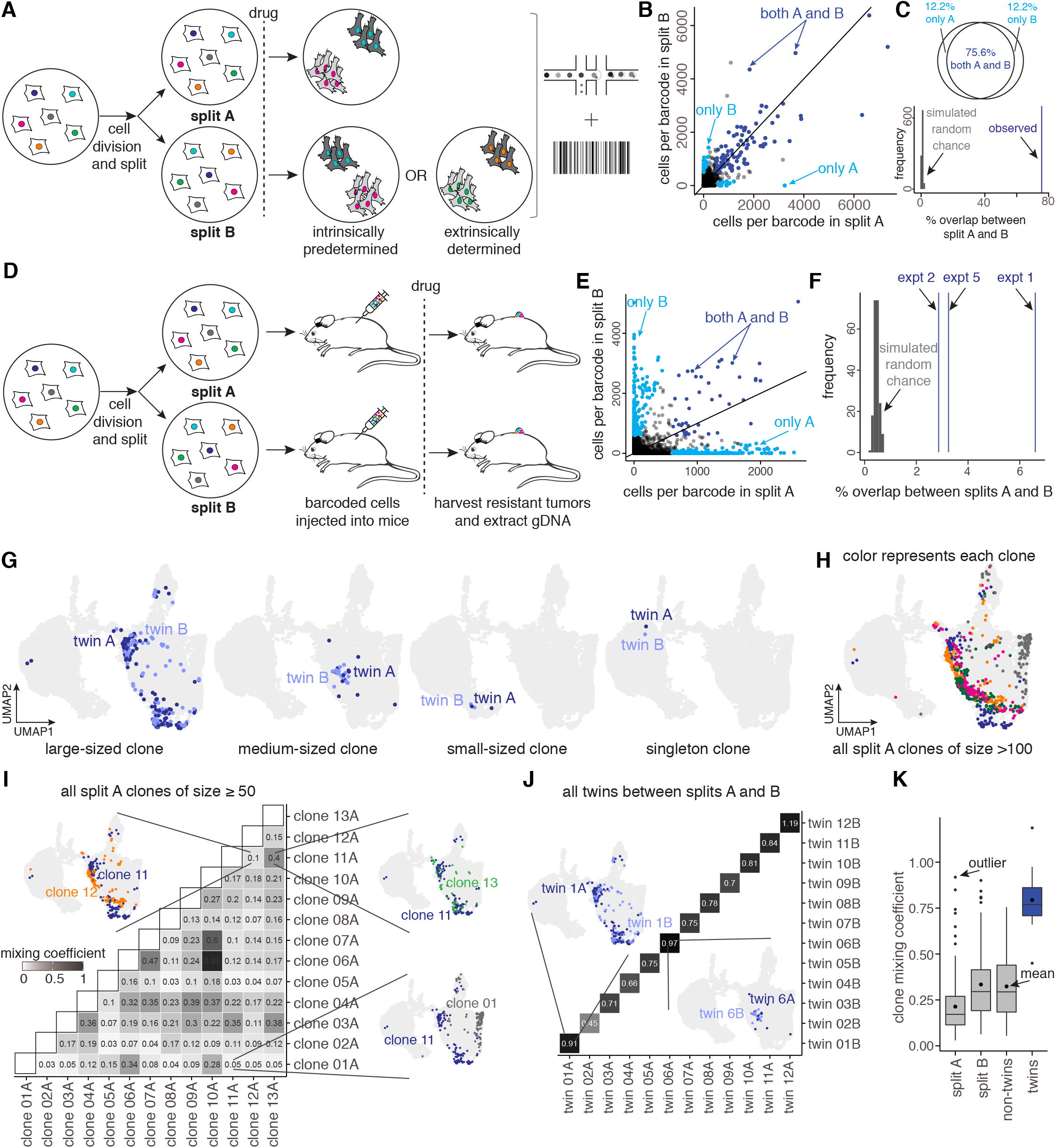
Cells are predestined for distinct resistant fates upon exposure to targeted therapy. a. Schematic of FateMap for following the fates of WM989 A6-G3 melanoma cells. For the experiment shown, we transduced WM989 A6-G3 cells at an MOI of ~0.15 with a barcode library similar to that described in (Emert et al., 2021). After 3-4 cell divisions, we sorted the successfully barcoded population (GFP positive), divided the culture in splits (A and B), treated each with vemurafenib, and performed single-cell RNA sequencing and barcode sequencing on the resultant colonies. b. For each barcode identified by sequencing of the genomic DNA, we plotted its abundance in corresponding splits A and B. Based on the observed strong correlation in barcode abundance between parallel cultures (splits A and B), we reasoned that the ability to survive and become resistant is strongly predetermined by the molecular state of the population of melanoma cells preceding drug exposure. Those present in both splits are colored in dark blue, and those present only in either A or B are colored in cyan. c. (top) A venn diagram showing a significant fraction of overlap between barcode clones present in both splits (dark blue) compared to those present only in either A or B (cyan). (bottom) Comparison of the observed overlap between the shared barcodes (twins) surviving across splits with a simulation for a random chance of survival (simulated 1,000 times). d. Schematic of experimental design for following the barcodes that survive and develop resistance in mice. For the experiment shown, we transduced WM989 A6-G3 with a barcode library similar to that described in (Emert et al., 2021) and after 4-5 divisions, we sorted the successfully barcoded population (GFP positive), divided the culture in splits (A and B), and injected them subcutaneously in different mice, which were then treated with a 417 mg PLX4720/kg diet. The resistant tumors were harvested and the barcode containing genomic DNA was simplified and sequenced for all mice. e. For each barcode identified by sequencing of the genomic DNA, we plotted its abundance in corresponding splits A and B. Those present in both splits are colored in dark blue, and those present only in either A or B are colored in cyan. f. Comparison of the observed overlap between the shared barcodes (twins) surviving across splits with a simulation for a random chance of survival (simulated 1,000 times). g. UMAPs of representative twin clones (sharing the same barcode) across the two splits A and B. The twins largely end up with the same transcriptional fate, invariant of the clone size. This observation suggests that cells are predestined for distinct resistant fates upon exposure to vemurafenib. h. Painting large clones onto the UMAP, where each color represents a unique resistant clone. i. We formulated a quantitative metric, the mixing coefficient, to calculate the transcriptional relatedness of different clones to each other. For each pair of barcoded clones, we calculated the nearest neighbors for each cell in the 50-dimensional principal component space. We then classified the neighbors as “self” if the neighbors are from the same barcode clone or “non-self” if they belong to another barcode clone. The mixing coefficient is the ratio of averaged fraction of self neighbors to averaged fraction non-self neighbors. The higher the mixing coefficient, the higher the transcriptional relatedness of the barcoded clones analyzed (a value of 1 corresponds to perfect mixing, and a value of 0 corresponds to no mixing). As shown with representative examples on the UMAP, for non-twin clones in split A, the mixing coefficient is largely low. j. The mixing coefficient for twin clones across splits A and B is presented with representative examples on the UMAP. k. Box-plots of comparison of cumulative mixing coefficients between clones within Split A and B, non-twin clones across A and B, and twin clones. This analysis revealed that the mixing coefficient for twins is much higher than non-twins, irrespective of the split.

To show that this predetermined ability to survive *in vitro* held *in vivo*, we injected barcoded and recently divided sets of “twins” subcutaneously into different mice then treated with a BRAF targeted therapy drug (Figure 4D). Upon extracting and sequencing barcodes from each of the resistant tumors, we found a significant percentage (4.24% +- 1.19% (SEM)) of barcodes appeared in resistant tumors from multiple mice. This overlap percentage was significantly more than what one would expect from a random chance of survival across the mice (0.46% +- 0.00% (SEM) (Figure 4E,F, see Methods), showing that the ability to survive and become resistant was also predetermined in mouse xenograft models.

Given that the ability to become resistant was predetermined, we next asked whether the specific fate that a resistant clone adopted was similarly predetermined by the initial state of the pre-resistant cell as opposed to external factors. That is, did twins separated into the two split populations (thus randomizing the position in the plate and neighboring cells) adopt similar or distinct transcriptional profiles after drug treatment? By comparing the transcriptional profiles of clones sharing the same barcode in the two split populations, we were able to compare the resistant fate outcomes. The initial inspection of the UMAP projections suggested that the recently divided twins surviving therapy largely end up in the same regions of the UMAP space (Figure 4G; Supplementary Figure 1J), and are more similar than non-twin barcodes belonging to similar fate clusters (Figure 4H).

To formalize the comparison of single-cell transcriptional profiles of clones across split populations, we formulated a metric we called the “mixing coefficient” (see Methods). The mixing coefficient provides a pairwise comparison of transcriptional similarity between any two clones in principal component space. Briefly, for each pair of clones (twin or non-twin), we measured, for each cell, the number of nearby neighbor cells (from the pair of clones) that were either from the same clone (self-neighbor) or the pair clone. We then averaged the fraction of self-neighbors across all cells from the clone pair and divided that by the averaged fraction of non-self neighbors and reported that as the mixing coefficient. A mixing coefficient of 1 signifies a high degree of shared transcriptionally similarity between the pair, while a mixing coefficient of 0 implies that the two clones are transcriptionally separated in the principal component space (see Methods). For unrelated (non-twin) clones, we observed a low mixing coefficient (Figure 4I; Supplementary Figure 5). However, the twins sharing the same barcode across the splits A and B exhibited high mixing coefficients (Fig. 4J), which were significantly higher than pairwise comparisons of non-twins either within or across the two split populations (Fig. 4K). The fact that twin cells adopt a similar resistant fate upon drug exposure suggests that the adoption of distinct transcriptional and phenotypic fates was predetermined by the intrinsic molecular state of the cells preceding drug exposure, and environmental factors had little to no effect on the fate outcomes.

We also tested whether resistant fate predestination occurred in another melanoma cell line (WM983B A6-G3) and the breast cancer cell line tested earlier (MDA-MB-231-D4) (Supplementary Figure 3,4). For WM983B E6-C6, when we performed a similar experiment separating twins across split populations, we found that twins exposed to vemurafenib treatment largely ended up with the same resistant fate (Supplementary Figure 3G), much like the main melanoma cell line we analyzed. Similarly, FateMap on the MDA-MB-231-D4 breast cancer cells confirmed these phenotypic correspondences (Shaffer et al., 2020), with multiple dominant resistant clones being present in each of the two split populations, again suggesting a strong tendency for twins sharing the same barcode to become resistant to paclitaxel, a cytotoxic chemotherapy (Supplementary Figure 4I). The three largest clones appeared to occupy similar transcriptional profiles (Supplementary Figure 4H), although the effect was not as strong as what we observed for the melanoma cells in that the clones were not as restricted in transcriptional space. (Another factor could be barcode silencing, which we observed in many of the resistant colonies from MDA-MB-231-D4 cells (GFP silencing in ~23% of colonies in a dish) (Supplementary Figure 4E), thus making them harder to detect by FateMap as FateMap relies on the barcode to be transcribed into the cytoplasm.)

### Changing the drug dose results in fate switching and altered transcriptional profiles of some resistant fates

Our results revealed that for a particular drug concentration, there is a mapping between the initial molecular state of cells preceding drug exposure to the drug-resistant fate they ultimately adopt. However, drug resistance is known to depend heavily on the concentration used. We wondered how the ensemble of resistant fates would change if we used a different drug concentration. In principle, there were many possibilities, including the fates staying the same, but with different relative proportions, or the emergence of completely new transcriptional fates. To determine which of these possibilities occurred, we employed a similar experimental design to the twin experiments above, except that we exposed the two split populations to different dosage of the drug.

We verified that 100nM (above the IC50 concentration) was a reasonable low dose of vemurafenib, which eliminated the Ras/ERK signaling in these cells (Supplementary Figure 6A). With 100nM vemurafenib as the low dose and 1μM vemurafenib as the high dose (the concentrations used throughout the rest of the paper), we found that the number of resistant clones emerging from treatment with the low dose were ~2.5 fold higher than that for the high dose, with the same number of starting cells (Figure 5A; Supplementary Movie 1,2). We saw a similar behavior in mice, where the resistant tumors grew comparatively faster in the low dose (Figure 5B).

**Figure 5:**
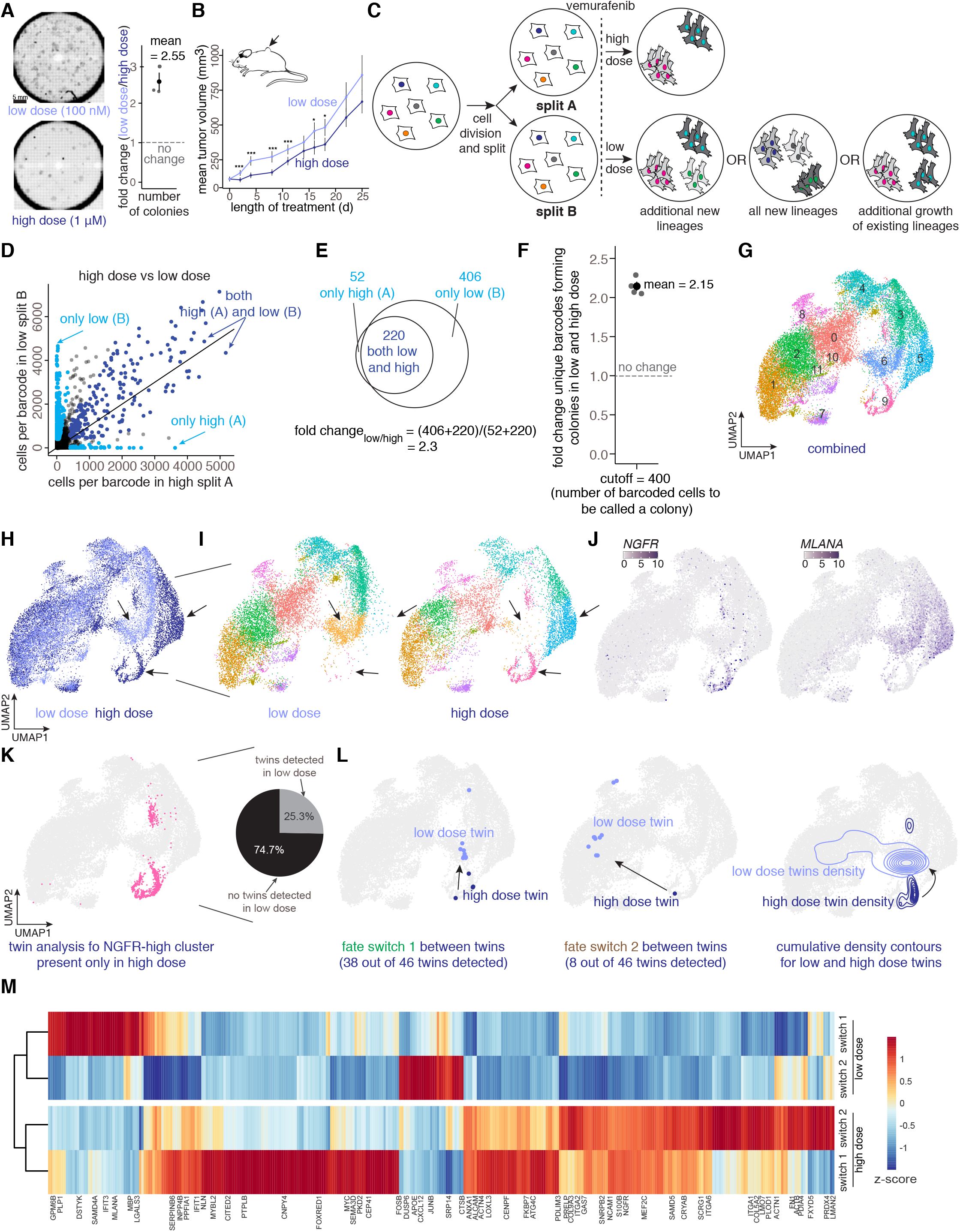
Changing the therapy dose results in stereotypic fate switching and altered transcriptional profiles of resistant fates. a. (left) Nuclei scans (DAPI-stained) of resistant colonies emerging from treatment of 25,000 WM989 A6-G3 cells with two different doses of vemurafenib (1uM and 100nM). (right) Quantification of the total number of colonies from each dose across biological replicates shows a 2-3 fold increase in colonies in the low dose as compared to the high dose. b. The resistant tumors grew faster early on in the low dose (41.7 mg PLX4720/kg) as compared to the high dose (417 mg PLX4720/kg). (t-test, *** = p < 0.05; * = 0.1 > p >0.05). c. Schematic of FateMap for following the fates of WM989 A6-G3 melanoma cells when exposed to two different doses of vemurafenib. For the experiment shown, we transduced WM989 A6-G3 cells at an MOI of ~0.15 with a barcode library similar to that described in Emert et al. 2021. After 3-4 cell divisions, we sorted the successfully barcoded population (GFP positive), divided the culture in splits (A and B), treated each with vemurafenib, and performed single-cell RNA sequencing and barcode sequencing on the resultant colonies. Given the increase in the number of colonies in the low dose, either of the three scenarios listed in the split B arm are possible. d. For each barcode identified by sequencing, we plotted its abundance in corresponding splits A (high dose) and B (low dose). Those present in both high and low dose splits are colored in dark blue, and those present only in either A (high) or B (low) are colored in cyan. Those present in both (dark blue) exhibited a strong correlation, suggesting that their ability to survive and become resistant is invariant of drug dose. For those present only in either (cyan), we found them to be much more abundant in low dose (split B), suggesting that new barcodes that are otherwise unable to survive in the high dose become drug-resistant in the low dose. e. A venn diagram showing that while there is a significant fraction of overlap between barcode clones present in both high and low dose splits (dark blue), those present in only low (B) comprise a significant fraction as well. f. Quantification of the total number of unique barcodes forming colonies from each dose across biological replicates shows a 2.15 average fold increase in colonies in the low dose as compared to the high dose. This ratio is close to what is observed from imaging experiments in A, suggesting that additional independent clones in the low dose alone can explain the increased number of colonies in the low dose. g. Combined low and high dose resistant cells obtained from UMAP applied to the first 50 principal components. Cells are colored by clusters determined using Seurat’s FindClusters command at a resolution of 0.8 (i.e. “Seurat clusters, resolution = 0.8”). h. UMAP where the resistant cells are colored by the associated dose; dark blue represents the high dose and light blue represents the low dose. Arrows represent UMAP regions that are present only in the low or high dose. i. UMAP is split by each dose, with colors representing clusters determined using Seurat’s FindClusters, and corresponding to those in g. Arrows represent UMAP regions that are present only in the low or high dose. j. UMAPs recolored for each cell by its expression for genes *NGFR* and *MLANA*, which are markers for clusters enriched in either of the two doses (as shown with arrows). k. (left) UMAP cluster colored for only cluster 9 (high for *NGFR*). (right) A pie chart to demonstrate that of all the clones (barcodes) present in high dose in cluster 9, 25.3% were also present in the low dose, albeit in different regions of the UMAP. l. (left) Two representative examples of UMAP regions where twins from the *NGFR*-high cluster in the high dose go to the low dose. Of all the twins present in both (46), 38 behave more similarly to each other than the second group (fate switch 1), while the other eight behave more similarly to each other than the first group (fate switch 2). (right) A cumulative density contour plot capturing the two types of fate switches that *NGFR*-high cluster clones from the high dose adopt in the low dose. m. A heatmap of differentially expressed genes for fate switch 1, fate switch 2, and between *NGFR*-high clones exhibiting fate switches 1 and 2.

The increase in the number of resistant clones at low dose could have arisen for a number of reasons. One possibility was that additional new clones became resistant besides those that survived in the case of high dose treatment (Figure 5C). Alternatively, completely different sets of clones may survive in low dose and have minimal overlap with those becoming resistant in the high dose (Figure 5C). Yet another possibility is that no new unique clones arise, and that the additional resistant clones in the low dose case are a result of additional divisions of the same set of pre-resistant cells in the low dose (Figure 5C). We used FateMap to distinguish between these possibilities.

Using an experimental design similar to the previous section, we barcoded cells, allowed them to divide, separated them into different groups, and this time exposed split population A to high dose and split population B to low dose (Figure 5C). We first analyzed the survival of clones in each of the high and low dose split populations after 3-4 weeks of treatment by sequencing the gDNA of the surviving populations. Similar to our results from treatment with the same dosage of drug, we found a significant overlap between unique barcodes detected in surviving cells in both drug concentrations (Figure 5D,E; Supplementary Figure 6F). However, we also observed many barcodes that only survived in low concentration, implying that additional new clones survive the low dose and reprogram to become resistant. The average fold-change increase in resistant clones in low dose from barcode sequencing (2.15) was strikingly similar to the fold-change increase from imaging (2.55) (Figure 5A,E-F; Supplementary Figure 6B).

We wondered if the clones that only survived at low dose were also predetermined to survive low dose. That is, if we exposed two separated twin split populations to the same low dose, would the clones that were detected only in low dose appear across low dose split populations? We found that the clones present in only the low dose also tended to show up across two low dose split populations, showing that these low-dose only clones are also the result of intrinsic pre-existing differences between initial cells as opposed to microenvironmental differences (Supplementary Figure 6C-E).

We next asked how the diversity of resistant clones changed between low and high doses of drug. Did new transcriptional fates arise at the lower dose? Did the proportions change between existing fates? Or some combination of the two? We performed a joint FateMap analysis of both the low and high dose resistant populations combined (Figure 5G). Broadly, despite the overall large changes in the nature of resistance between low and high dose, we found that many of the transcriptional fates were the same between the two doses (Figure 5H-J; Supplementary Figure 6G). However, there were many significant differences as well. Particularly, the *NGFR*-high cells (cluster 9) were largely missing from the low dose resistant population of cells. Additionally, while the *MLANA*-high cells (clusters 5,6) were present at both doses, they were in non-overlapping clusters (Figure 5H-J).

Given these differences, we asked whether the twins corresponding to the *NGFR*-high cells at high dose existed at low dose, and if so, what resistant fate they adopted. We collected all the barcoded-clones in the high dose corresponding to the *NGFR*-high cells (cluster 9) and looked for their corresponding twins in the low dose (Figure 5K). We found that 46 such barcodes (25.3% of all barcodes in cluster 9) were indeed present in the low dose (Figure 5K). This number is much higher than what one would expect from random chance (Figure 4C, bottom). When we analyzed the fates of these barcodes in the low dose, we found that a vast majority (38/46) of the clones adopted fates within the *MLANA*-high cluster 6 (fate switch 1) (Figure 5L). The remaining clones appeared to adopt a different, albeit less transcriptionally constrained fate (fate switch 2) (Figure 5L). These observations, presented with two examples, are captured by the cumulative density contours for all 46 shared twins between low and high dose (Figure 5L). We found several genes to be differentially expressed for pairwise comparisons of the fate switches as compared to the corresponding twins in the high dose NGFR cluster (149 genes for switch 1 and 216 genes for switch 2) (Figure 5M; Supplementary Table 8).

We wondered whether there were subtle differences between cells within the high-dose *NGFR* positive cluster that indicated whether the twin cells at low dose adopted the *MLANA*-high fate or not. We found that within the high dose, when we compared the 38 clones with the remaining 8 clones, we found 70 genes to be differentially expressed, a gene number higher than comparing randomly sampled cells from *NGFR*-high cluster 9 (p-value <0.05) (Figure 5M; Supplementary Figure 6H), suggesting that subtle pre-existing differences between cells that appeared to adopt the same *NGFR*-high fate at high dose could lead to more obvious fate differences at low dose.

For the *MLANA*-high resistant fate clusters, we found that while this fate largely consisted of singletons (4.66% non-singletons clones) in the high dose, this fate contained a far higher fraction of non-singleton clones in the low dose (21.6% non-singletons) (Figure 5J; Supplementary Figure 6I). Thus, we observed a change in phenotype (increased proliferation) of resistant fate of cells between high and low dose.

### Changing the therapy type results in fate switching and altered transcriptional profiles of resistant fates

We next wondered what happens to the ensemble of fates when we expose the cells to different inhibitors of MAPK signaling. To answer this, we used a FateMap design similar to previous sections and exposed the split populations A and B to BRAF inhibitor vemurafenib (1μM) and MEK inhibitor trametinib (5nM), respectively. We found again that while many resistant cells from each drug have similar transcriptional profiles (Figure 6A), there were significant differences as well (Figure 6A,B). Of particular interest, we saw a depletion of *MLANA*-high cells (cluster 3) in trametinib compared to vemurafenib (Figure 6B,F). Cells from this cluster largely existed as singletons, i.e. non-colony forming resistant cells, at this dose of vemurafenib, so we wondered if trametinib treatment resulted in fewer singleton cells in general. Looking at cluster 3, we saw that the number of singletons were significantly higher in vemurafenib compared to trametinib (Figure 6C). To confirm the sequencing-based results, we used imaging to verify the reduction in the number of singletons (Figure 6D; Supplementary Movie 1,3), which revealed that while the number of colonies was similar in each condition (fold change = 1.23), the singletons, were about 3-fold higher in vemurafenib compared to trametinib (Figure 6D,E). This observation shows that trametinib, a MEK inhibitor, selectively inhibits the formation of the singleton cell fate i.e. *MLANA*-high cells (Figure 6F). Most of the vemurafenib cluster 3 clones (95.2% of all clones) did not have corresponding twins in trametinib (Figure 6G), suggesting that those cells were actually killed by trametinib as opposed to just being converted to a different resistant fate, in stark contrast to the fate of a missing cluster (*NGFR*-high cells) when comparing across low and high dose of vemurafenib (25.3% of twin clones survived survived across doses) (Figure 5K). Those few clones present (~4.8%) tend to largely adopt a fate in the similar UMAP regions (clusters 3 and 4), although some do end up adopting a completely different fate (Figure 6H-I; Supplementary Figure 7A).

**Figure 6:**
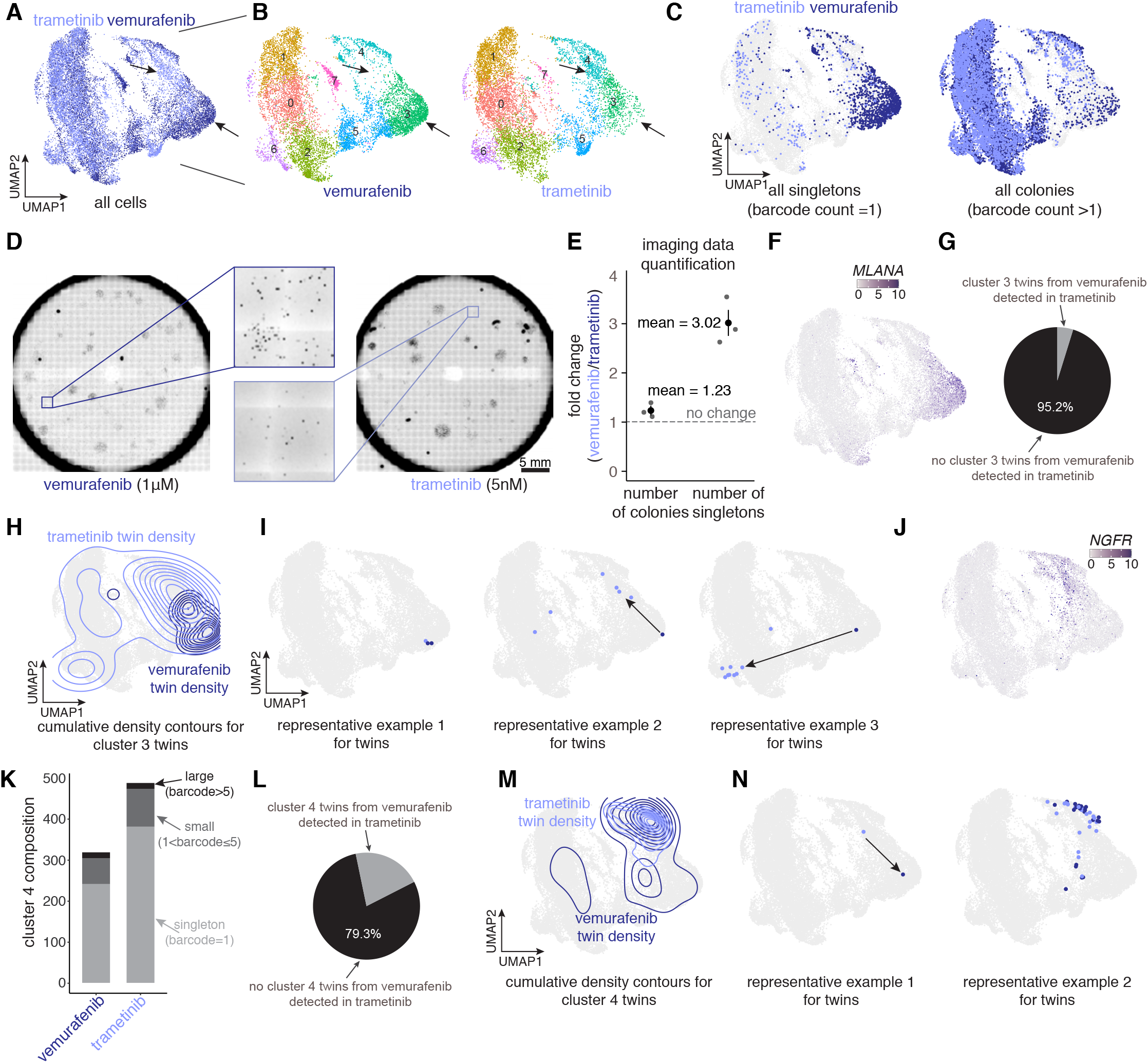
Changing the therapy type to trametinib eliminates an additional resistant fate present in the vemurafenib treatment. a. UMAP where the resistant cells are colored by the associated therapy drug type, with dark blue representing vemurafenib and light blue representing trametinib. Arrows represent UMAP regions that are present only in vemurafenib or trametinib. b. UMAP is split by each drug type, with colors representing clusters determined using Seurat’s FindClusters command at a resolution of 0.5 (i.e. “Seurat clusters, resolution = 0.5”). Arrows represent UMAP regions that are present only in vemurafenib or trametinib. c. Painting of singletons and colonies onto the UMAP, colored by the condition, demonstrated that singletons largely belong to vemurafenib and are present predominantly in the *MLANA*-high cluster. Colonies are dispersed more across the UMAP with no particular region enriched for either condition except for the *NGFR*-high cluster. d. Imaging of nuclei (DAPI-stained) of resistant colonies emerging from treatment of WM989 A6-G3 cells to either vemurafenib or trametinib. The number of singletons in trametinib treated cells appear to be much less than those treated with vemurafenib, consistent with the sequencing data from FateMap. e. Quantification of the total number of colonies and singletons from each drug type of imaging data across biological replicates. This analysis demonstrated that while the total number of colonies are similar across the two drug types, there is a relative increase in the number of singletons in the case of vemurafenib. f. UMAPs are recolored for each cell by its expression for gene *MLANA*, which is a marker for cluster 3 relatively enriched in vemurafenib (as shown with arrows in A and B). g. A pie chart to demonstrate that of all the clones (barcodes) present in vemurafenib-treated split in cluster 3, only 4.8% were also present in the trametinib-treated split. h. A cumulative density contour plot capturing the types of fate switches that the *MLANA*-high cluster 3 clones from the vemurafenib-treated split adopt in the trametinib-treated split. i. Three representative examples of UMAP regions where twins from the *MLANA*-high cluster 3 in the vemurafenib-treated split adopt in the trametinib-treated split. j. UMAPs are recolored for each cell by its expression for gene *NGFR*, which is a marker for cluster 4 relatively enriched in trametinib (as shown with arrows in A and B). k. Composition of clones of different sizes within *NGFR*-high cluster 4 for both trametinib- and vemurafenib-treated splits. l. A pie chart to demonstrate that of all the clones (barcodes) present in the trametinib-treated split in cluster 4, 20.7% were also present in the vemurafenib-treated split. m. A cumulative density contour plot capturing the types of fate switches that the *NGFR*-high cluster 4 clones from the vemurafenib-treated split adopt in the trametinib-treated split. n. Two representative examples of UMAP regions where twins from the *NGFR*-high cluster 4 in trametinib-treated split adopt in the vemurafenib-treated split.

With FateMap, we also found the *NGFR*-high cluster 4 to be relatively more populated in the trametinib-treated cells as compared to vemurafenib treatment (Figure 6B,J,K), which we verified by imaging (Supplementary Figure 7B,C). Unlike the *MLANA*-high cells in cluster 3, for which FateMap clone barcodes appearing in the vemurafenib-treated split population were not generally found in the trametinib-treated population(Figure 6G), we found that for the *NGFR*-high cluster 4, a much higher percentage of barcodes from the trametinib-treated split population (20.7%) were also present in the vemurafenib-treated split population. The twins of these trametinib-treated *NGFR*-high cells adopted either the same fate (*NGFR*-high) or in some cases *MLANA*-high fate in vemurafenib (Figure 6L-N). Therefore, changing the type of drug can significantly affect the nature and proportions of drug-resistant fates.

We also wondered if the dual treatment of vemurafenib and trametinib may kill off additional resistant fates or if the dual treatment mimics treatment with one of the two drugs alone. We found that the transcriptional profiles of resistant fates in dual treatment are virtually indistinguishable from that of trametinib alone (Supplementary Figure 7D,E). This overlap suggests that, at least for the doses tested, trametinib has a dominant role in guiding fate outcomes.

### Inhibition of histone methyltransferase DOT1L increases the number of resistant colonies with largely the same fates

We next wondered how the fate outcomes are affected upon altering the intrinsic molecular state of cells prior to exposing them to therapies. Genetic screening from our lab identified DOT1L, a histone methyltransferase, as a modulator of single-cell variability (Torre et al., 2021). Inhibition of DOT1L prior to application of the drug increasing the frequency of resistant clones. We asked whether the additional resistant colonies appearing with DOT1L inhibition adopted new fates than without inhibition.

We first compared the survival of resistant clones between DOT1L inhibitor-pretreated and DMSO-pretreated (control) conditions. We found that many more barcoded clones survived vemurafenib treatment following pretreatment with a DOT1L inhibitor (pinometostat) compared to the control (DMSO) (Figure 7A) (Emert et al., 2021). Therefore, increased resistance in DOT1L inhibitor condition is at least partially due to the survival of barcoded clones that otherwise do not survive in the control condition.

**Figure 7:**
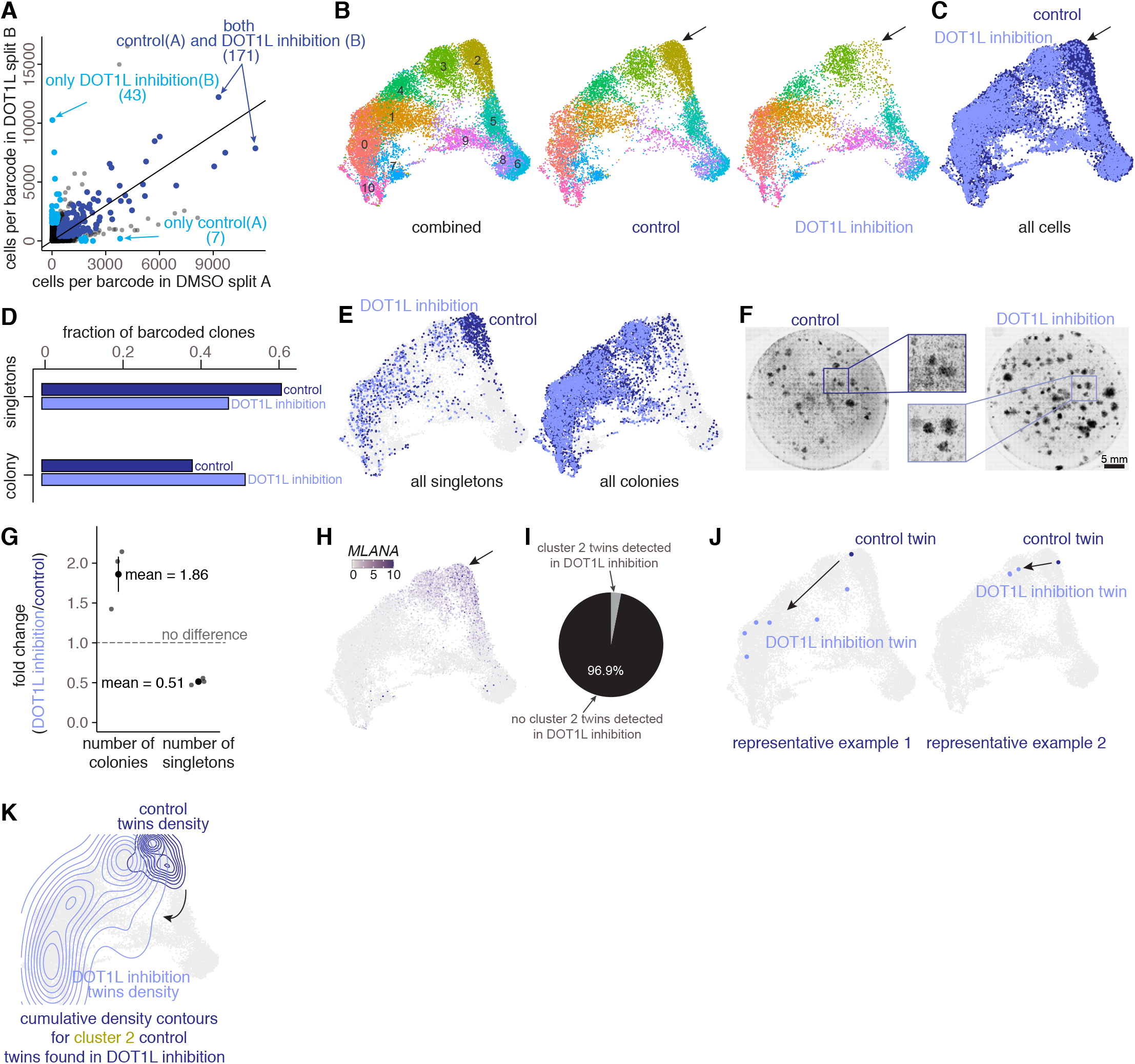
Inhibition of histone methyltransferase DOT1L results in the emergence of additional resistant proliferative clones and a reduction in singletons. a. For each barcode identified by sequencing, we plotted its abundance in corresponding splits A (DMSO control) and B (DOT1L inhibition). Those present in both control and DOT1L splits are colored in dark blue, and those present only in either A (control) and B (DOT1L) are colored in cyan. Those present in both (dark blue) exhibited a strong correlation, suggesting that their ability to survive and become resistant is invariant of drug dose. For those present only in either (cyan), we found them to be much more abundant in DOT1L (B), suggesting that new barcodes, otherwise unable to survive in the control condition, become drug-resistant in the DOT1L inhibited condition. b. (left) Combined resistant cells in the control and DOT1L conditions obtained from UMAP applied to the first 50 principal components. Cells are colored by clusters determined using Seurat’s FindClusters command at a resolution of 0.8 (i.e. “Seurat clusters, resolution = 0.8”). (right) UMAP is split by each condition. c. UMAP where the resistant cells are colored by the associated condition (control vs DOT1L); dark blue represents control and light blue represents DOT1L. The arrow represents the UMAP region present predominantly in the control region and missing from the DOT1L-associated UMAP region. d. Quantification of singletons and colonies showed that while the number of resistant colonies is higher in DOT1L, it is accompanied by a reduced number of singletons cells compared to control. e. Painting of singletons and colonies onto the UMAP, colored by the condition, demonstrated that singletons largely belong to the control condition and are present predominantly in cluster 2 (*MLANA*-high). Colonies are dispersed more across the UMAP with no particular region enriched for either condition. f. Imaging of the nuclei (DAPI-stained) of resistant colonies emerging from vemurafenib treatment of WM989 A6-G3 cells, either for control or cells lacking DOT1L. g. Quantification of the total number of colonies and singletons from each type across biological replicates demonstrated a relative increase in total colonies and reduction in total singletons in the DOT1L and control conditions, respectively. h. UMAP is recolored for each cell by its expression for the gene *MLANA*, a marker for cluster 2, which is relatively enriched in control (as shown with an arrow). i. A pie chart to demonstrate that of all the clones (barcodes) present in the control condition split, only 3.1% were also present in the DOT1L inhibitor pretreatment split. j. Two representative examples of UMAP regions where twins from the *MLANA*-high cluster in the control condition go in the DOT1L condition. A cumulative density contour plot capturing the types of fate switches that *MLANA*-high cluster clones from control adopt in the DOT1L inhibitor-pretreated condition. k. A cumulative density contour plot capturing the types of fate switches that the *MLANA*-high cluster 2 clones from the control condition split adopt in the DOT1L inhibitor pretreatment split.

Upon performing FateMap, we found that DOT1L inhibition prior to treatment with vemurafenib did not change most of the resultant resistant fates (Figure 7B,C). However, the *MLANA*-high transcriptional cluster (Figure 7H), corresponding to largely singletons in control, had a significantly smaller number of resistant cells from DOT1L inhibitor-pretreated cells (Figure 7D,E). The resistant colonies that emerged from the DOT1L inhibitor-pretreatment cells adopted resistant fates similar to the control condition (Figure 7E, Supplementary Figure 7F), in contrast to what we observed when comparing low dose to high dose (Figure 5). We confirmed these observations by direct imaging (Figure 7F,G).

Given that the *MLANA*-high cells are depleted in DOT1L inhibitor-pretreated cells (cluster 2), we wondered what fates did cluster 2 twins in control, if present, adopt in the DOT1L inhibitor-pretreated condition (Figure 7H). We collected all the barcoded clones in *MLANA*-high cluster 2 and found that a majority of them (96.9%) are not present in the DOT1Li-pretreated condition, suggesting that these singletons largely don’t survive the DOT1Li-pretreated condition (Figure 7I). Those present (3.1%) tended to span the entirety of the transcriptional space (Figure 7J,K), unlike the case with altering drug dose, where we observed a clear fate switch.

## Discussion

Much work has gone into the consequences of single-cell variability on responses to external cues, but little has been done to systematically quantify variability in the outcomes in otherwise homogeneous populations. Here, we used FateMap to reveal extensive variability in the outcome of cells after the cue, in this case between resistant cancer cells after treatment with targeted therapies. These outcomes are largely predetermined by molecular differences in the initial state of cells, some of which have been elucidated (Emert et al., 2021). What these results suggest is that even homogeneous-seeming cells have a large amount of latent variability that can be revealed by various cues. We suspect that cues that cause cellular stress may, in particular, allow this variability to express itself by stabilizing and enhancing the preexisting differences, akin to removing evolutionary “capacitors” (Queitsch et al., 2002; Rutherford and Lindquist, 1998; Waddington, 2014).

Such results suggest the possibility of a rich mapping between the initial molecular states of cells and their outcome. We have further shown that these mappings are strongly dependent on the nature of the external cue—different doses and drugs can lead to important differences in terms of which cells adopt what fates. Hence, the perturbation itself must be considered an integral part of the mapping, and the mappings between initial state and ultimate fate are not meaningful without specification of the external cue.

Two important factors that underlie cell fate determination in response to a cue are the memory of the initial state and the degree to which extrinsic factors influence the ultimate fate determination. Our system exhibited enough memory (meaning persistence of the state through multiple divisions (Emert et al., 2021; Shaffer et al., 2020)) to allow us to perform twin experiments, and then those twins largely adopted the same fates, showing that in this context, ultimate cell fate was largely intrinsically determined. This intrinsic determination need not always be the case. For instance, performing a similar analysis on cardiac differentiation, we did not observe predetermination of fate upon exposure of hiPS cells to differentiation signals (Jiang et al., 2021). Instead, in that case, cell fate was largely determined by extrinsic factors. It is also possible to have a situation in which there is short memory, but fate determination is still largely intrinsic. Such cases would be hard to discriminate because twin experiments would show little correspondence in the fates of twins, even though the state of the twins before the cue is given is still largely determining the outcome.

An important therapeutic implication of our work is that different treatments can lead to subtle but important differences in drug-resistant fates. For instance, we showed that treatment with trametinib, a MEK inhibitor, was able to eliminate specific resistant fates that otherwise continue to persist in treatment with the canonical drug vemurafenib, a BRAF inhibitor (both of which impinge on the same molecular pathway (Goyal et al., 2017)). These differences may help explain the effectiveness of newer dual therapy regimens combining both MEK and BRAF inhibition in melanoma patients (Robert et al., 2019).

It remains unclear what regulatory processes are at play in determining the ultimate fates. One view is that there is a complex set of specific regulatory interactions that create certain “attractor states” that are a consequence of the cell’s existing intrinsic regulatory program. Supporting this view is the fact that we observed many of the same resistant fates across a number of drug doses and combinations. An alternative possibility is that cells undergo a general adaptation to whatever stress they are faced with, enabling a wider range of outcomes than the cell’s default regulatory program may allow. The fact that we observe differences between virtually every resistant clone (but twins are largely identical) suggests that there is some general adaptive process at work that “burns in” all intrinsic differences in the initial state, even minor ones, regardless of their functional consequence. Furthermore, the finding that each individual resistant clone seemed to have different expression profiles, even when belonging to the same broad transcriptional class, also suggests an adaptive response (Figure 4H). These two types of fate determination would have conceptually distinct mechanisms underlying them; future work would do well to try and decipher these mechanisms.

We mapped morphology and functional properties, e.g. invasion, by comparing bulk sequencing of manually isolated and expanded resistant clones to single-cell transcriptomic data from FateMap which do not necessarily guarantee purity or equivalent comparisons. Future work can modify the FateMap framework to directly measure functional properties, such as invasion or interaction with vasculature, by performing these assays on barcoded cells in well-controlled microengineered devices (Park et al., 2019; Wouters et al., 2020). In a similar vein, while much of our work has focused on molecular and functional characterization of the diverse fates individually in 2D tissue culture systems, it remains unclear how the emerging variability manifests in 3D. FateMap, in conjunction with *in situ* imaging of barcoded populations (Feldman et al., 2019), can be leveraged to reveal how the resistant fates interact with each other and more generally the ecological principles underlying tumor progression.

Here, we focused on characterizing resistant fates of single-cell-derived cancer cells in the face of anti-cancer therapies. Since non-genetic variability underlies several biological processes, including stem cell reprogramming and directed differentiation (Jiang et al., 2021; Pour et al., 2015), FateMap could be used in these contexts to reveal the gamut of emergent fates and identify their potential origins.

## Methods

### Cell lines and culture

WM989 A6-G3 and WM983b E9-C6 melanoma cell lines, first described in (Emert et al., 2021) and kindly provided by the lab of Dr. Meenhard Herlyn, were derived by twice single-cell bottlenecking the WM989 and WM983b melanoma cell lines respectively. The identities of WM989 A6-G3 and WM983B E9-C6 were verified (Emert et al., 2021) by DNA STR microsatellite fingerprinting at the Wistar Institute. WM989 A6-G3 5a3, first described in (Torre et al., 2021), was derived by single-cell bottlenecking of WM989 A6-G3. MDA-MB-231-D4, first described in (Shaffer et al., 2020) was derived by single-cell bottlenecking of MDA-MB-231 (ATCC HTB-26). The identity of MDA-MB-231-D4 was verified (Shaffer et al., 2020) by ATCC human STR profiling cell line authentication services.

WM989 A6-G3, WM989 A6-G3 5a3, and WM983b E9-C6 melanoma cell lines were cultured in TU2% media (80% MCDB 153, 10% Leibovitz’s L-15, 2% FBS, 2.4mM CaCl2, 50 U/mL penicillin, and 50 μg/mL streptomycin). MDA-MB-231 cell lines were cultured in DMEM10% (DMEM with glutamax, 10% FBS and 50 U/mL penicillin, and 50 μg/mL streptomycin). All three cell lines were passaged using 0.05% trypsin-EDTA.

### Flow sorting of barcoded cells

We used 0.05% trypsin-EDTA (Gibco, 25300120) to detach the barcoded cells from the plate and subsequently neutralized the trypsin with the corresponding media depending on the cell type (Tu2% for WM989 and WM983B; DMEM + 10% FBS for MDA-MB-231). We then pelleted the cells, performed a wash with 1X DPBS (Invitrogen, cat. 14190-136), and resuspended them again in 1X DPBS. Cells were sorted on a BD FACSJazz machine (BD Biosciences), gated for positive GFP signal and singlets. Sorted cells were then centrifuged to remove the supernatant media containing PBS, and replated with the appropriate cell culture media.

### Drug treatment experiments

We prepared stock solutions in DMSO of 4 mM vemurafenib (PLX4032, Selleck Chemicals, S1267), 10 mM pinometostat (Selleck Chemicals, S7062), 100 μM trametinib (Selleck Chemicals, S2673), and 4mM paclitaxel (Life Technologies, P3456). We prepared small aliquots (10-15ul) for each drug and stored them at −20 °C to minimize freeze–thaw cycles. For drug treatment experiments, we diluted the stock solutions in culture medium to a final concentration of 1μM and 100nM for vemurafenib, 4 μM for pinometostat, 5 nM for trametinib, and 1 nM for paclitaxel unless otherwise specified.

WM989 A6-G3 and WM983b E9-C6 cells were treated with either vemurafenib or trametinib for 3-4 weeks, and the media was replaced every 3-4d. Similarly, MDA-MB-231-D4 cells were treated with paclitaxel for 3-4 weeks, and the media was replaced every 3-4d. At the end of the treatment, surviving cells were trypsinized, neutralized, washed with 1X DPBS, and then either 1) pelleted and stored at −20°C for gDNA extraction, or 2) resuspended in PBS for single-cell RNA sequencing experiments. In some cases, cells were also fixed for imaging at the end of the treatment. For pinometostat (DOT1L inhibitor) pre-treatment (before addition of vemurafenib), WM989 A6-G3 cells were treated for five days, replacing media once at day 3.

### RNA FISH on cells in plates

We performed single-molecule RNA FISH as previously described (Raj et al., 2008). For the genes used in this study, we designed complementary oligonucleotide probe sets using custom probe design software (MATLAB) and ordered them with a primary amine group on the 3’ end from Biosearch Technologies (Supplementary Table 3 for probe sequences). We then pooled each gene’s complementary oligos and coupled the set to Cy3 (GE Healthcare), Alexa Fluor 594 (Life Technologies) or Atto647N (ATTO-TEC) N-hydroxysuccinimide ester dyes.

The cells were fixed as follows: we aspirated media from the plates containing cells, washed the cells once with 1X DPBS, and then incubated the cells in the fixation buffer (3.7% formaldehyde in 1X DPBS) for 10 min at room temperature. We then aspirated the fixation buffer, washed samples twice with 1X DPBS, and added 70% ethanol before storing samples at 4°C. For hybridization of RNA FISH probes, we rinsed samples with wash buffer (10% formamide in 2X SSC) before adding hybridization buffer (10% formamide and 10% dextran sulfate in 2X SSC) with standard concentrations of RNA FISH probes and incubating samples overnight with coverslips, in humidified containers at 37°C. The next morning, we performed two 30-min washes at 37°C with the wash buffer, after which we added 2X SSC with 50 ng/mL of DAPI. We mounted the sample(s) for imaging in 2X SSC.

### Immunofluorescence and imaging

For NGFR staining of fixed cells, after fixation and permeabilization, we washed the cells for 10 min with 0.1% BSA/PBS, and then stained the cells for 30 min with 1:500 anti-NGFR APC-labeled clone ME20.4 (BioLegend, 345107). We washed the cells five times with 0.1% BSA/PBS and followed with one final wash with PBS for 2 min at room temperature. Fresh PBS was added prior to imaging. Wells were imaged either immediately or after storage in 4°C overnight. All conditions (wells) were fixed, permeabilized, and stained at the same time with identical settings. Wells from the same plate were all imaged consecutively in the same imaging session.

For dpERK staining of fixed cells, after fixation and permeabilization, we used primary antibodies targeting dpERK (p44/p42 ERK D12.14.4E Cell Signaling, 4370). First, we rinsed cells twice for 5 min each time with 5% BSA in PBS (5% BSA-PBS) and then incubated in the dark at room temperature for 2 hours in 5% BSA-PBS 1:200 dpERK antibodies. Next, we washed the cells 5 X 5 min with 5% BSA-PBS and then incubated the cells at room temperature for 1 hour in 5% BSA-PBS containing 1:500 goat anti-rabbit secondary antibody conjugated to Alexa Fluor 594 (Cell Signaling, 8889). After the secondary incubation, we washed the cells 5 X 5 min with 5% BSA-PBS containing 50 ng/mL of DAPI and then replaced the wash buffer with fresh PBS and proceeded with imaging consecutively. All conditions (wells) were fixed, permeabilized, stained, and imaged at the same time with identical settings.

For colony counting via nuclei imaging, the cells were fixed by aspirating media from the plates containing cells, washing the cells once with 1X DPBS, and then incubating the cells in the fixation buffer (3.7% formaldehyde in 1X DPBS) for 10 min at room temperature. We aspirated the fixation buffer, washed samples twice with 1X DPBS, and added 70% ethanol before storing samples at 4°C. Fixed cells were stained for nuclei by incubation in 2X SSC containing 50 ng/ml of DAPI and then imaged each well *via* a tiling scan at 10X magnification.

### Barcode lentivirus library generation and diversity estimation

Barcode libraries were constructed as previously described (Emert et al., 2021), and the protocol is available at https://www.protocols.io/view/barcode-plasmid-library-cloning-4hggt3w. Briefly, we modified the LRG2.1T plasmid (gift from Dr. Junwei Shi) by removing the U6 promoter and single guide RNA scaffold. We then inserted a spacer sequence flanked by EcoRV restriction sites after the stop codon of GFP, subsequently digesting this vector backbone with EcoRV (NEB) and gel purifying the linearized vector. We ordered PAGE-purified ultramer oligonucleotides (IDT) containing 100 nucleotides with a repeating “WSN” pattern (W = A or T, S = G or C, N = any) surrounded by 30 nucleotides homologous to the vector insertion site (Supplementary Table 1). We subsequently used Gibson assembly followed by column purification to combine the linearized vector and barcode oligo insert. We performed nine electroporations of the column-purified plasmid into Endura electrocompetent *Escherichia coli* cells (Lucigen) using a Gene Pulser Xcell (Bio-Rad). We then allowed for their recovery before plating serial dilutions and seeding cultures for maxi-preparation. We incubated these cultures on a shaker at 225 rpm and 32°C for 12-14 hours, pelleted the resulting cultures by centrifugation, and used the EndoFree Plasmid Maxi Kit (Qiagen) to isolate plasmid according to the manufacturer’s protocol. Barcode insertion was verified by polymerase chain reaction (PCR) on colonies from plated serial dilutions. We pooled the plasmids from the 9 separate cultures in equal amounts by weight before packaging into lentivirus.

To estimate the barcode library complexity, we performed three independent transductions (see below for details) on WM989 A6-G3 melanoma cell lines, extracted gDNA, sequenced the barcodes, and noted the total and overlapping barcodes between pairs of three independent transductions. We estimated the barcode library complexity with the equation used in mark and capture analysis: k/K = n/N, where k is number of recaptured barcodes that were marked, K is number of barcodes captured in the second pool, n is the number of barcodes marked in the first pool, N is the estimated barcode library complexity. Using this formula, we found the barcode diversity from three transductions to be 48.9, 54.4, and 63.3 million barcodes (Supplementary Figure 1C; raw data and calculation scripts are within the link provided in Data and Code Availability).

### Lentivirus packaging and transduction

We adapted previously described protocols to package lentivirus (Emert et al., 2021; Torre et al., 2021). We first grew HEK293FT to near confluency (80-95%) in 10cm plates in DMEM containing 10% FBS and 50 U/mL penicillin, and 50 μg/mL streptomycin, and one day before plasmid transfection, we changed the media in HEK293FT cells to DMEM containing 10% FBS without antibiotics. For each 10cm plate, we added 80 μL of polyethylenimine (Polysciences, cat. 23966) to 500 μL of Opti-MEM (Thermo Fisher Scientific, cat. 31985062), separately combining 5 μg of VSVG and 7.5 μg of pPAX2 and 7.35 μg of the barcode plasmid library in 500 μL of Opti-MEM. We then incubated both solutions separately at room temperature for 5 min. We then mixed both solutions together by vortexing and incubated the combined plasmid-polyethylenimine solution at room temperature for 15 min. We added 1.09 mL of the combined plasmid-polyethylenimine solution dropwise to each 10cm dish. After 6-7 hours, we aspirated the media from the cells, washed the cells with 1X DPBS, and added fresh TU2% media. The next morning, we aspirated the media, and added fresh Tu2% media. Approximately 9-11 hours later, we transferred the virus-laden media to an empty, sterile 50ml tube and stored it at 4C, and added fresh Tu2% media to each plate. We continued to collect the virus-laden media every 9-11 hours for the next ~30 hours in the same 50ml tube, and stored the collected media at 4C. Upon final collection, we filtered the virus-laden media through a 0.45μm PES filter (MilliporeSigma SE1M003M00) and stored 1.5ml aliquots in cryovials at −80°C.

To transduce WM989 A6-G3, WM983b E9-C6, and MDA-MB-231-D4 cells, we freshly thawed virus-laden media on ice, added it to dissociated cells, and plated ~100,000 cells/well in a six-well plate with ~3ml of the media. We then centrifuged the 6-well plate at 1,750 r.p.m. (517g) for 25 min. We then incubated the 6-well plate at 37°C and replaced the media at ~8h, washed with 1X DPBS, and added fresh media (TU2% for WM989 and WM983B, and DMEM with 10% FBS for MDA-MB-231) to each well. After ~24 hours, we passaged the cells to 10-cm dishes. The barcoded cells (GFP-positive) were then sorted and plated for a total of 4-5 population doubling until treatment with appropriate drugs. The time to 4-5 population doubling was 10-11 days for WM989, 6-7 days for WM983B, 5-6 days for MDA-MB-231. The volume of the virus-laden media was decided by the titers performed on each cell line and target multiplicity of infection (MOI). For single-cell RNA sequencing experiments in particular, we targeted for the MOI to be ~10-25% to minimize the fraction of cells with multiple unique barcodes. We found it to be relatively computationally challenging to differentiate multiple-barcoded cells from doublets introduced by gel beads-in-emulsions.

### Single-cell RNA sequencing

We used the 10X Genomics single-cell RNA-seq kit v3 to sequence barcoded cells. We resuspended the cells (targeting ~10,000 cells for recovery/ sample) in PBS and followed the protocol for the Chromium Next GEM Single Cell 3’ Reagent Kits v3.1 as per manufacturer directions (10X Genomics, Pleasanton, CA). Briefly, we generated gel beads-in-emulsion (GEMs) using the 10X Chromium system, and subsequently extracted and amplified (11 cycles) barcoded cDNA as per post-GEM RT-cleanup instructions. We then used a fraction of this amplified cDNA (25%) and proceeded with fragmentation, end-repair, poly A-tailing, adapter ligation, and 10X sample indexing per the manufacturer’s protocol. We quantified libraries using the High Sensitivity dsDNA kit (Thermo Fisher Q32854) on Qubit 2.0 Fluorometer (Thermo Fisher Q32866) and Bioanalyzer 2100 (Agilent G2939BA) analysis prior to sequencing on a NextSeq 500 machine (Illumina) using 28 cycles for read 1, 55 cycles for read 2, and 8 cycles for i7 index.

### Computational analyses of single-cell RNA sequencing expression data

We adapted the cellranger v3.0.2 by 10X Genomics into our custom pipeline (https://github.com/arjunrajlaboratory/10XCellranger) to map and align the reads from NextSeq sequencing run(s). Briefly, we downloaded the bcl counts and used *cellranger mkfastq* to demultiplex raw base call files into library-specific FASTQ files. We aligned the FASTQ files to the hg19 human reference genome and extracted gene expression count matrices using *cellranger count*, while also filtering and correcting cell identifiers and unique molecular identifiers (UMI) with default settings.

We then performed the downstream single-cell expression analysis in Seurat v3. Within each experimental sample, we removed genes that were present in less than three cells, as well as cells with less than or equal to 200 genes. We also filtered for mitochondrial gene fraction which was dependent on the cell type. For non-identically treated samples, we integrated them using scanorama (Hie et al., 2019), which may work better to integrate non-similar datasets and avoid over-clustering. For samples that were exposed to identical treatment, we normalized using SCTransform (Hafemeister and Satija, 2019) and the samples according to the Satija lab’s integration workflow (https://satijalab.org/seurat/articles/integration_introduction.html). Using scanorama on identically-treated samples produced qualitatively similar results (Supplementary Figure 1K,L).

For each experiment, we used these integrated datasets to generate data dimensionality reductions by principal component analysis (PCA) and Uniform Manifold Approximation and Projection (UMAP), using 50 principal components for UMAP generation. For a majority of analyses, we worked with the principal component space and normalized expression counts. For rare cases where we used Seurat UMAP clusters, we tested a range of resolutions with Seurat’s FindClusters command. Our conclusions did not change qualitatively when we tested resolutions between 0.4 and 1.2 (Supplementary Figure 2).

### Bulk sequencing and analysis

We conducted standard bulk paired-end (37:8:8:38) RNA sequencing using RNeasy Micro (Qiagen 74004) for RNA extraction, NEBNext Poly(A) mRNA Magnetic Isolation Module (NEB E7490L), NEBNext Ultra II RNA Library Prep Kit for Illumina (NEB E7770L), NEBNext Multiplex Oligos for Illumina (Dual Index Primers Set 1) oligos (NEB E7600S), and an Illumina NextSeq 550 75 cycle high-output kit (Illumina 20024906), as previously described (Mellis et al., 2021; Shaffer et al., 2017). Prior to extraction and library preparation, the samples were randomized to avoid any experimental and human biases. As previously described, we aligned RNA-seq reads to the human genome (hg19) with STAR v2.5.2a and counted uniquely mapping reads with HTSeq v0.6.1 (Dobin et al., 2013; Mellis et al., 2021; Shaffer et al., 2017) and outputs count matrix. The counts matrix was used to obtain tpm and other normalized values for each gene using scripts provided at: https://github.com/arjunrajlaboratory/RajLabSeqTools/tree/master/LocalComputerScripts

To compare bulk sequencing data with single-cell RNA sequencing datasets, we first extracted differentially expressed genes for each single-cell RNA sequencing cluster (snn = 0.6) using the Seurat command FindAllMarkers, and filtering for adjusted p-value <0.05 and avg_logFC > 1. Similarly, we extracted differentially expressed genes for each condition of interest (morphology or invasiveness) and filtering for −1.5 < avg_logFC > 1.5. We then calculated the similarity score, which represents the normalized fraction of overlap of differentially expressed genes for the condition of interest between bulk-sequencing data and each single-cell RNA sequencing cluster.

### Expanded resistant colonies morphology categorization

The resistant colonies were manually binned in one of the three categories based on the morphology images taken from the Nikon TS2-FL microscope: “small”, “on top”, and “not on top”. Those that were difficult to be binned in any category were labeled as “uncategorized”. Of the three categories, the “small” category was the most easy to identify manually and label due to characteristic optical and proliferation (slow growing) properties. The other two categories (“on top” and “not on top”) had further sets of morphological and proliferative differences, but were difficult to be parsed into specific categories. Of the 64 resistant colonies isolated and expanded across therapy treatments of vemurafenib and trametinib, five were uncategorized. The differentially upregulated genes for “small” and “on top” are provided in Supplementary Table 4. For category “not on top”, only four genes were differentially upregulated, thus precluding us from doing further analysis.

### Nearest neighbor analysis

We developed a quantifiable approach to measure the gene expression relatedness of different barcoded clones. For each pair of barcoded clones, we calculated the nearest neighbors for each cell in the 50-dimensional principal component space. We then classified the neighbors as “self” if the neighbors are from the same barcode clone or “non-self” if they belong to the other barcode clone. We defined a quantifiable metric, the mixing coefficient, as follows:

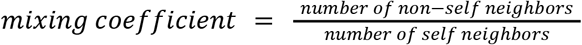

A mixing coefficient of 1 would indicate perfect mixing such that each cell has the same number of self and non-self neighbors. A mixing coefficient of 0 would indicate that there is no mixing and that each cell within a barcoded clone lies far away from the other barcoded clone in the principal component space. The higher the mixing coefficient, the higher the transcriptional relatedness of the barcoded clones analyzed. As the number of nearest neighbors depends on the size (number of cells) of a clone, we performed this analysis between cells of similar clone size (Supplementary Figure 5). Within a specified size range, we normalized the number of neighbors per barcode clone to account for small size differences. The number of neighbors to extract was chosen to be a minimum of 10 and the size of the smaller of the two barcode clones.

### Barcode recovery from single-cell RNA sequencing data

As the barcodes are transcribed, we extracted the barcode information from the amplified cDNA from 10X Genomics V3 chemistry protocol (step 2). We ran a PCR side reaction with one primer that targets the 3’ UTR of GFP and the other that targets a region introduced by the amplification step within the V3 chemistry of 10X genomics (“Read 1”). The two primers amplify both the 10X cell-identifying sequence as well as the 100 bp barcode that we introduced lentivirally. The number of cycles, typically between 12-15, are decided by the Ct value from a qPCR reaction (NEBNext Q5 Hot Start HiFi PCR Master Mix (New England Biolabs)) for the specified cDNA concentration. The thermal cycler (Veriti 4375305) was set to the following settings: 98 °C for 30 s, followed by N cycles of 98 °C for 10 s and then 65 °C for 2 min and, finally, 65 °C for 5 min. Upon completion of the PCR reaction, we immediately performed a 0.7X bead purification (Beckman Coulter SPRISelect) followed by final elution in nuclease-free water. Purified libraries were quantified with High Sensitivity dsDNA kit (Thermo Fisher) on Qubit Fluorometer (Thermo Fisher), pooled, and sequenced on a NextSeq 500. We sequence 26 cycles on Read 1 which gives 10X cell-identifying sequence and UMI, 124 cycles for read 2 which gives the barcode sequence, and 8 cycles for index i7 to demultiplex pooled samples. The primers used are provided in Supplementary Table 2.

### Experiments in mice

For each experiment, WM989 cells were uniquely barcoded with protocols as described above and allowed to divide 4-5 times before splitting the barcoded pool into five groups, each containing an equal number of cells. We aimed for ~1-1.5 million WM989 cells to be injected per mice. All animal experiments were performed in accordance with institutional and national guidelines and regulations. The protocols have been approved by the Wistar IACUC. WM989 cells in serum-free RPMI 1640 media (Corning 10-40-CM) were mixed in a 1:1 ratio with Growth Factor Reduced Matrigel® (Corning 354230), then were subcutaneously implanted into the flanks of NSG mice. Once tumors reached about 100mm^3^ per caliper measurement, animals were randomized into treatment groups. Treatment consisted of Low Dose 41.7 mg PLX4720/kg diet (Research Diets D21051202i), or High Dose 417 mg PLX4720/kg diet (Research Diets D21051201i), to which they had constant access. Tumor size was measured by calipers every 2-4 days, and tumor volumes were calculated according to the equation 0.5*L*W*W, where L is the longest side and W is a line perpendicular to L. Mice were sacrificed once tumors reached 1500 mm^3^, and once one mouse reached the endpoint, all mice from the same barcode pool were sacrificed regardless of tumor size. The tumor tissue was snap frozen in liquid N_2_ for genomic DNA extraction. We had 5 biological replicate experiments. We could not extract sufficient gDNA from experiments 3 and 4, and these were excluded from barcode split population analysis.

### Computational analyses of barcoded single-cell datasets

The barcodes from the side reaction of single-cell cDNA libraries were recovered by developing custom shell, R, and python scripts, which are all available at this link: https://github.com/arjunrajlaboratory/10XBarcodeMatching. Briefly, we scan through each read searching for sequences complementary to the side reaction library preparation primers, filtering out reads that lack the GFP barcode sequence, have too many repeated nucleotides, or do not meet a phred score cutoff. Since small differences in otherwise identical barcodes can be introduced due to sequencing and/or PCR errors, we merged highly similar barcode sequences using STARCODE software (Zorita et al., 2015), available at https://github.com/gui11aume/starcode. For varying lengths of barcodes (30, 40 or 50, see the pipeline guide provided) depending on the initial distribution of Levenshtein distance of non-merged barcodes, we merged sequences with Levenshtein distance ≤ 8, summed the counts, and kept only the most abundant barcode sequence.

For next processing steps and downstream analysis, we first filtered out all barcodes that were associated below the minimum cutoff (dependent on sequencing depth) of unique molecular identifiers (UMI). We next removed all barcodes where one 10X cell-identifying sequence was associated with more than one unique barcode. This could either result from multiplets introduced within gel beads-in-emulsions or because of the same cell receiving multiple barcodes during lentiviral transduction. After these two filtering steps, we were able to recover barcodes associated with 50-60% of single cells, which were then used to do the downstream clone-resolved analysis.

### Barcode library preparation and sequencing from genomic DNA

We prepared barcode libraries from genomic DNA (gDNA) as previously described (Emert et al., 2021). Briefly, we isolated gDNA from barcoded cells using the QIAmp DNA Mini Kit (Qiagen, cat. 51304) per the manufacturer’s protocol. Extracted gDNA was stored as a pellet in −20°C for days to weeks before the next step. We then performed targeted amplification of the barcode using custom primers containing Illumina adaptor sequences, unique sample indices, variable-length staggered bases, and an ‘UMI’ consisting of 6 random nucleotides (NHNNNN). As reported in (Emert et al., 2021), the ‘UMI’ does not uniquely tag barcode DNA molecules, but nevertheless appeared to increase reproducibility and normalize raw read counts. We determined the number of amplification cycles (N) by initially performing a separate quantitative PCR (qPCR) and selecting the number of cycles needed to achieve one-third of the maximum fluorescence intensity for serial dilutions of genomic DNA. The thermal cycler (Veriti 4375786) was set to the following settings: 98°C for 30 s, followed by N cycles of 98°C for 10 s and then 65°C for 40 s and, finally, 65 °C for 5 min. Upon completion of the PCR reaction, we immediately performed a 0.7X bead purification (Beckman Coulter SPRISelect), followed by final elution in nuclease-free water. Purified libraries were quantified with a High Sensitivity dsDNA kit (Thermo Fisher) on a Qubit Fluorometer (Thermo Fisher), pooled, and sequenced on a NextSeq 500 using 150 cycles for read 1 and eight cycles for each index (i5 and i7). The primers used are provided in Supplementary Table 6.

### Analyses of sequenced barcodes from genomic DNA

The barcode libraries from genomic DNA sequencing data were analysed as previously described (Emert et al., 2021), with the custom barcode analysis pipeline available at https://github.com/arjunrajlaboratory/timemachine. Briefly, this pipeline searches for barcode sequences that satisfy a minimum phred score and a minimum length. Note that we count the total number of ‘UMIs’ as described in the gDNA library chemistry above. These ‘UMIs’ do not necessarily tag unique barcode DNA molecules, but empirically they slightly improve correlation in barcode abundance among replicate libraries (Emert et al., 2021). We also use STARCODE (Zorita et al., 2015), available at https://github.com/gui11aume/starcode, to merge sequences with Levenshtein distance ≤ 8 and add the counts across collapsed (merged) barcode sequences.

In this current work, we also created two subclones (D8 and F8) of WM989 A6-G3, with each clone carrying a unique barcode sequence (Supplementary Table 7). We used these two clones as standards to convert sequencing counts into actual cell numbers which significantly reduces the PCR and cell number bias across samples. We spiked in a known number of cells from each of the two barcoded clones to each cell pellet before gDNA extraction and sequencing. We then used linear regression (on (0,0), (count_F8, cells_F8), (count_D8, cells_D8)) to get the conversion factor from read counts of all barcodes to their actual cell numbers. We used a minimum cell count and log2-fold change between pairs of conditions to annotate clones as condition-dependent or condition-independent. We found that changing the cutoff for minimum cell count did not affect our conclusions (Supplementary Figure 6B).

### Simulation for barcode overlap

We adapted a described previously computational model that simulates all steps of our experiments designed to compare barcode overlap in resistant colonies (Yunusova et al., 2017). The model simulates cell seeding and infection. Each cell is represented as an independent object. The number of barcoded cells was calculated as

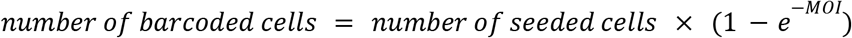

where the MOI was estimated for our barcode lentivirus. Barcodes were represented by integer numbers from among 50 million variants of unique barcodes estimated from our lentiviral library diversity (see Methods and Supplementary Figure 1C). The subset of barcoded cells was assigned barcodes randomly with replacement from this library. The model simulates expanding cells prior to addition of the drug. Each cell, regardless of barcode status, undergoes a cell division procedure with 4-5 rounds depending on the experimental condition. In each round, a given cell will give rise to a number of progeny sharing the same barcode based on an estimated distribution of cell division. The model plates cells onto separate dishes/splits (total dishes/splits dependent on the experiment) by randomly assigning each cell an integer. The model simulates the formation of resistant colonies assuming a purely stochastic model of resistance. A defined fraction of cells on each plate form resistant colonies based on an resistance efficiency that was calculated as

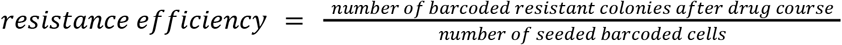

based on experimental observations. Additionally, each cell forming a resistant colony is subject to a probabilistic material loss at different stages of the *in silico* experiment, including cell culture (5% both *in vivo* and *in vitro*), genomic DNA extraction (0% *in vitro* and 10% *in vivo* mouse), DNA sequencing library preparation (5% both *in vivo* and *in vitro*), and (as needed) mouse injection (15%) and tumor extraction (15%). The output of the model was the number of barcodes shared between different plates or barcode overlap. This was not corrected for cells having more than one lentiviral barcode due to multiple integrations for a given MOI. We performed 1000 and 200 independent simulations for *in vitro* and *in vivo* experiments respectively to obtain a distribution of barcode overlap values to determine the probability of obtaining our observed barcode overlap from our experiments by random chance. This model was written and executed in R.

### Tissue sectioning and RNA FISH

We adapted the protocol described previously (Symmons et al., 2019). Tumor tissue extracted from mice subcutaneously injected with WM989 A6-G3 5a3 (Torre et al., 2021) was mounted in Tissue-Plus O.C.T. compound (Fisher Healthcare), flash-frozen in liquid nitrogen, wrapped in aluminium foil, and then stored tissues at −80°C. Tissues were cryosectioned at 6 or 8 μm using a Leica CM1950 cryostat within the Center for Musculoskeletal Disorders (PCMD) Histology Core. We adhered tissue samples to positively charged Superfrost Plus slides (Fisher Scientific). We then washed slides in PBS, fixed them in 4% formaldehyde for 10 min at room temperature, then washed them two times in PBS. Fixed slides were stored in 70% ethanol in LockMailer microscope slide jars at 4°C.

For RNA FISH on tissue sections, we placed the slide in the wash buffer (2X SSC, 10% formamide) and allowed it to equilibrate for 2-3 min. We then removed slides from the wash buffer and dried off the slides with kimwipes. Immediately after drying, we added 500-1000ul of 8% SDS to the tissue section on the slide for 1 min. After 1 min, we turned the slide on the side to remove the SDS, transferred the slide to the wash buffer, and kept it in the wash buffer for ~2 min. We then tapped down the wash buffer on a kimwipe or paper towel, added 50 μL of probe-containing hybridization buffer (10% dextran sulfate, 2X SSC, 10% formamide) as a drop in center of tissue sample, and placed a cover slip on top of the tissue section. We then placed the slide into a humidifying chamber to prevent the slide from drying, and placed the chamber containing slides in a 37°C incubator overnight. We took out the chamber to RT the next day and placed the slide with coverslip into a wash buffer container and let the cover slip come off. We then transferred the slides to a container or LockMailer jar containing wash buffer and incubated for 30 min at 37°C. We removed it from the 37°C incubator and performed a second incubation with wash buffer and DAPI, and put it back into 37°C for another 30 min. We performed one final wash in wash buffer, rinsed two times in 2X SSC, and added 50-100uL of 2X SSC to the tissue section. We then placed a coverslip on the tissue, sealed it with nail varnish, and let it dry before imaging.

### Spheroid assay

We adapted the protocol described previously (Kaur et al., 2019). Tissue culture–treated 96-well plates were coated with 50 μL 1.5% Difco Agar Noble (Becton Dickinson). Melanoma cells were seeded at 3000 cells/well and allowed to form spheroids over 96 to 120 hours. Spheroids were harvested and embedded using collagen type I (GIBCO, #A1048301). The collagen plug was prepared as 300 μL mix per layer, and two layers were added into each well [1X Eagle Minimum Essential Medium (EMEM; 12-684, Lonza); 10% FCS; 1X l-glutamine; 1.0 mg/mL collagen I; NaHCO3 (17-613E, Lonza), diluted in PBS as required]. The first layer was added to each well and allowed to solidify. After 5 to 10 min, spheroids were mixed with the remaining 300 μL mix and added to the well to solidify. Once the plug was solidified, media were added to the well and incubated at 37°C at 5% CO2 and imaged after 24 hours and 48 hours. Spheroid images were acquired on a Nikon Ti2E inverted microscope. Quantitation of invasive surface area was performed using NIS Elements Advanced Research software. Of the 64 resistant colonies we expanded, only 24 colonies had enough cells to form multiple spheroids per colony. Of these 24, 6 did not form spheroids and 2 only had one spheroid each. Of the remaining 16 colonies, 8 colonies belonged to resistant colonies emerging from trametinib, 3 belonged to 1μM vemurafenib and 5 belonged to 250nM vemurafenib. The differentially upregulated genes for “fast invading” and “slow invading” are provided in Supplementary Table 5.

### Imaging

To image RNA FISH and nuclei signal, we used a Nikon TI-E inverted fluorescence microscope equipped with a SOLA SE U-nIR light engine (Lumencor), a Hamamatsu ORCA-Flash 4.0 V3 sCMOS camera, and 4X Plan-Fluor DL 4XF (Nikon MRH20041/MRH20045), 10X Plan-Fluor 10X/0.30 (Nikon MRH10101) and 60X Plan-Apo λ (MRD01605) objectives. We used the following filter sets to acquire different fluorescence channels: 31000v2 (Chroma) for DAPI, 41028 (Chroma) for Atto 488, SP102v1 (Chroma) for Cy3, 17 SP104v2 (Chroma) for Atto 647N, and a custom filter set for Alexa 594. We tuned the exposure times depending on the dyes used (Cy3, Atto 647N, Alexa 594, and DAPI). For large tiled scans, we used a Nikon Perfect Focus system to maintain focus across the imaging area. For imaging RNA FISH signals in tissue sections, we acquired z-stacks (three positions) at 60X magnification, and used maximum intensity projection to visualize the signal. For brightfield imaging of resistant colonies, we used a Nikon Eclipse Ts2-FL with an Imagingsource DFK 33UX252 camera and 4X Plan-Fluor 4X/0.13 (Nikon MRH20041) objective. For time-lapse imaging of the emergence of drug-resistant colonies, we used an IncuCyte S3 Live Cell Imaging Analysis System (Sartorius) with a 4X objective on WM989 A6-G3 tagged with an mCherry nuclear reporter (H2B-mCherry).

### Image processing

For colony counting, all image processing was done blind to the condition (either drug type/dose or with/without DOT1L inhibition). The wells (within the 6-well plates) were pseudo-named in a format independent from drug or dose. Nikon-generated nd2 files were first parsed using custom MATLAB scripts (rajlabformattools) to convert them from nd2 format to tiff format. The code can be found here: https://github.com/arjunrajlaboratory/rajlabformattools. Images for each well were then stitched using custom MATLAB code and the number of cells in each well was counted using custom MATLAB code with a Gaussian filter consistent across samples being compared (colonycounting_v2). Colonies within each well were manually segmented and MATLAB was used to calculate the total number of colonies, cells per colony, and cells outside of colonies. The code can be found here: https://github.com/arjunrajlaboratory/colonycounting_v2.

For immunofluorescence, all image processing was done blind to drug conditions; the wells were pseudo-named in a format independent from drug or dose. Nikon images were first stitched using Nikon Elements software. The channels were split in Fiji and scaled to a smaller size compared to their original pixel size prior to placing them in illustrator. For the conditions being compared (e.g. each well with a different drug dose/type), the individual channels were equally adjusted for brightness and contrast across each pair of wells for the signal of interest. Raw nd2 files are provided for each imaging experiment which contains additional metadata for image settings. For the images taken on the brightfield microscope Nikon TS2FL, the scale bar lengths were calculated by using the pixel size given by the manufacturer of the camera. For well images taken on the Nikon TI-E inverted fluorescence microscope, the scale bar lengths were calculated using Nikon Elements software to add a line of a specific length to the images.

### Estimation of survival fraction in MDA-MB-231-D4

We estimated the frequency of drug resistance in MDA-MB-231-D4 by computing the fraction of surviving barcoded colonies upon treatment with paclitaxel as compared to the total number of uniquely barcoded cells in the initial population. From two separate split population experiments, we obtained the frequency to be 1:956 and 1: 1303 (see Data and Code Availability for the script).

## Data and Code Availability

All raw and processed data as well as code for the analyses in this manuscript can be found at https://www.dropbox.com/sh/my0vgswgrh03li4/AAC9R70cQwixeL-zYGsQumaka?dl=0.

## Author Contributions

YG and AR conceived and designed the project. YG designed, performed, and analyzed all experiments, supervised by AR. IPD assisted YG with tissue sectioning and automated RNA FISH and DAPI scans. YG, BE, and KK designed and optimized the PCR “side reaction” primers for recovering the barcodes from single-cell RNA sequencing libraries. AK assisted YG in the design and implementation of spheroid experiments. GTB, NJ, JL, and IAM assisted YG with barcode library preparation and computational pipeline. YG designed the mouse barcoding experiments, and DF and MEF performed the mouse experiments with input from YG, MH, AR, and ATW. YG and GTB prepared barcode libraries for mouse experiments. YG and AR wrote the manuscript with input from all authors.

## Acknowledgements

We thank Shweta Ramdas, Connie Jiang, Lauren Beck, Philip Burnham, Lee Richman, Samuel Reffsin, Eduardo Torre, Chris Cote, Allison Cote, Vito Rebecca, and Margaret Dunagin for insightful discussions related to this work. We thank the Genomics Facility at the Wistar Institute, especially Sonali Majumdar and Sandy Widura, for assistance with sequencing and single-cell partitioning and addition of 10X cell identifiers. We thank the Flow Cytometry Core Laboratory at the Children’s Hospital of Philadelphia Research Institute for assistance with flow cytometry and fluorescence-activated cell sorting. MEF and ATW thank the Core Facilities of the Johns Hopkins Kimmel Cancer Center, P30CA00697356. YG acknowledges support from the Burroughs Wellcome Fund Career Awards at the Scientific Interface, the Jane Coffin Childs Memorial Fund, and the Schmidt Science Fellowship; BE acknowledges support from NIH F30 CA236129, NIH T32 GM007170, and NIH T32 HG000046; AK acknowledges support from NIH K00-CA-212437-02; NJ acknowledges support from NIH F30 HD103378; IAM acknowledges support from NIH F30 NS100595; KK acknowledges support from NIH T32 GM008216; DF and MH acknowledge support from NIH grants RO1 CA238237, U54 CA224070, PO1 CA114046, P50CA174523 and the Dr. Miriam and Sheldon G. Adelson Medical Research Foundation; MEF and ATW acknowledge support from R01CA174746 and R01CA207935; ATW is also supported by a Team Science Award from the Melanoma Research Alliance and P01 CA114046; and AR acknowledges support from NIH Director’s Transformative Research Award R01 GM137425, NIH R01 CA238237, NIH R01 CA232256, NIH P30 CA016520, NIH SPORE P50 CA174523, and NIH U01 CA227550.

## References

Bhang, H.-E.C., Ruddy, D.A., Krishnamurthy Radhakrishna, V., Caushi, J.X., Zhao, R., Hims, M.M., Singh, A.P., Kao, I., Rakiec, D., Shaw, P., et al. (2015). Studying clonal dynamics in response to cancer therapy using high-complexity barcoding. Nat. Med. 21, 440–448.

Biddy, B.A., Kong, W., Kamimoto, K., Guo, C., Waye, S.E., Sun, T., and Morris, S.A. (2018). Single-cell mapping of lineage and identity in direct reprogramming. Nature 564, 219–224.

Chen, H., and Larson, D.R. (2016). What have single-molecule studies taught us about gene expression? Genes Dev. 30, 1796–1810.

Dobin, A., Davis, C.A., Schlesinger, F., Drenkow, J., Zaleski, C., Jha, S., Batut, P., Chaisson, M., and Gingeras, T.R. (2013). STAR: ultrafast universal RNA-seq aligner. Bioinformatics 29, 15–21.

Elowitz, M.B., Levine, A.J., Siggia, E.D., and Swain, P.S. (2002). Stochastic gene expression in a single cell. Science 297, 1183–1186.

Emert, B.L., Cote, C.J., Torre, E.A., Dardani, I.P., Jiang, C.L., Jain, N., Shaffer, S.M., and Raj, A. (2021). Variability within rare cell states enables multiple paths toward drug resistance. Nat. Biotechnol. 39, 865–876.

Feldman, D., Singh, A., Schmid-Burgk, J.L., Carlson, R.J., Mezger, A., Garrity, A.J., Zhang, F., and Blainey, P.C. (2019). Optical Pooled Screens in Human Cells. Cell 179, 787–799.e17.

Frieda, K.L., Linton, J.M., Hormoz, S., Choi, J., Chow, K.-H.K., Singer, Z.S., Budde, M.W., Elowitz, M.B., and Cai, L. (2017). Synthetic recording and in situ readout of lineage information in single cells. Nature 541, 107–111.

Goyal, Y., Jindal, G.A., Pelliccia, J.L., Yamaya, K., Yeung, E., Futran, A.S., Burdine, R.D., Schüpbach, T., and Shvartsman, S.Y. (2017). Divergent effects of intrinsically active MEK variants on developmental Ras signaling. Nat. Genet. 49, 465–469.

Gupta, P.B., Fillmore, C.M., Jiang, G., Shapira, S.D., Tao, K., Kuperwasser, C., and Lander, E.S. (2011). Stochastic state transitions give rise to phenotypic equilibrium in populations of cancer cells. Cell 146, 633–644.

Gutierrez, C., Al’Khafaji, A.M., Brenner, E., Johnson, K.E., Gohil, S.H., Lin, Z., Knisbacher, B.A., Durrett, R.E., Li, S., Parvin, S., et al. (2021). Multifunctional barcoding with ClonMapper enables high-resolution study of clonal dynamics during tumor evolution and treatment. Nature Cancer 2, 758–772.

Hafemeister, C., and Satija, R. (2019). Normalization and variance stabilization of single-cell RNA-seq data using regularized negative binomial regression. Genome Biol. 20, 296.

Hie, B., Bryson, B., and Berger, B. (2019). Efficient integration of heterogeneous single-cell transcriptomes using Scanorama. Nat. Biotechnol. 37, 685–691.

Jiang, C.L., Goyal, Y., Jain, N., Wang, Q., Truitt, R.E., Coté, A.J., Emert, B., Mellis, I.A., Kiani, K., Yang, W., et al. (2021). Cell type determination for cardiac differentiation occurs soon after seeding of human induced pluripotent stem cells.

Kaur, A., Ecker, B.L., Douglass, S.M., Kugel, C.H., 3rd, Webster, M.R., Almeida, F.V., Somasundaram, R., Hayden, J., Ban, E., Ahmadzadeh, H., et al. (2019). Remodeling of the Collagen Matrix in Aging Skin Promotes Melanoma Metastasis and Affects Immune Cell Motility. Cancer Discov. 9, 64–81.

Kinker, G.S., Greenwald, A.C., Tal, R., Orlova, Z., Cuoco, M.S., McFarland, J.M., Warren, A., Rodman, C., Roth, J.A., Bender, S.A., et al. (2020). Pan-cancer single-cell RNA-seq identifies recurring programs of cellular heterogeneity. Nat. Genet. 52, 1208–1218.

Krepler, C., Sproesser, K., Brafford, P., Beqiri, M., Garman, B., Xiao, M., Shannan, B., Watters, A., Perego, M., Zhang, G., et al. (2017). A Comprehensive Patient-Derived Xenograft Collection Representing the Heterogeneity of Melanoma. Cell Rep. 21, 1953–1967.

Leighton, J., Hu, M., Sei, E., Meric-Bernstam, F., and Navin, N.E. (2021). Reconstructing mutational lineages in breast cancer by multi-patient-targeted single cell DNA sequencing.

Marin-Bejar, O., Rogiers, A., Dewaele, M., Femel, J., Karras, P., Pozniak, J., Bervoets, G., Van Raemdonck, N., Pedri, D., Swings, T., et al. (2021). Evolutionary predictability of genetic versus nongenetic resistance to anticancer drugs in melanoma. Cancer Cell 39, 1135–1149.e8.

Mellis, I.A., Edelstein, H.I., Truitt, R., Goyal, Y., Beck, L.E., Symmons, O., Dunagin, M.C., Linares Saldana, R.A., Shah, P.P., Pérez-Bermejo, J.A., et al. (2021). Responsiveness to perturbations is a hallmark of transcription factors that maintain cell identity in vitro. Cell Syst 12, 885–899.e8.

Oren, Y., Tsabar, M., Cuoco, M.S., Amir-Zilberstein, L., Cabanos, H.F., Hütter, J.-C., Hu, B., Thakore, P.I., Tabaka, M., Fulco, C.P., et al. (2021). Cycling cancer persister cells arise from lineages with distinct programs. Nature 596, 576–582.

Park, S.E., Georgescu, A., Oh, J.M., Kwon, K.W., and Huh, D. (2019). Polydopamine-Based Interfacial Engineering of Extracellular Matrix Hydrogels for the Construction and Long-Term Maintenance of Living Three-Dimensional Tissues. ACS Appl. Mater. Interfaces 11, 23919–23925.

Pillai, M., and Jolly, M.K. (2021). Systems-level network modeling deciphers the master regulators of phenotypic plasticity and heterogeneity in melanoma. iScience 24.

Pour, M., Pilzer, I., Rosner, R., Smith, Z.D., Meissner, A., and Nachman, I. (2015). Epigenetic predisposition to reprogramming fates in somatic cells. EMBO Rep. 16, 370–378.

Queitsch, C., Sangster, T.A., and Lindquist, S. (2002). Hsp90 as a capacitor of phenotypic variation. Nature 417, 618–624.

Raj, A., and van Oudenaarden, A. (2008). Nature, nurture, or chance: stochastic gene expression and its consequences. Cell 135, 216–226.

Raj, A., Peskin, C.S., Tranchina, D., Vargas, D.Y., and Tyagi, S. (2006). Stochastic mRNA synthesis in mammalian cells. PLoS Biol. 4, e309.

Raj, A., van den Bogaard, P., Rifkin, S.A., van Oudenaarden, A., and Tyagi, S. (2008). Imaging individual mRNA molecules using multiple singly labeled probes. Nat. Methods 5, 877–879.

Rambow, F., Rogiers, A., Marin-Bejar, O., Aibar, S., Femel, J., Dewaele, M., Karras, P., Brown, D., Chang, Y.H., Debiec-Rychter, M., et al. (2018). Toward Minimal Residual Disease-Directed Therapy in Melanoma. Cell 174, 843–855.e19.

Ramirez, M., Rajaram, S., Steininger, R.J., Osipchuk, D., Roth, M.A., Morinishi, L.S., Evans, L., Ji, W., Hsu, C.-H., Thurley, K., et al. (2016). Diverse drug-resistance mechanisms can emerge from drug-tolerant cancer persister cells. Nat. Commun. 7, 10690.

Robert, C., Grob, J.J., Stroyakovskiy, D., Karaszewska, B., Hauschild, A., Levchenko, E., Chiarion Sileni, V., Schachter, J., Garbe, C., Bondarenko, I., et al. (2019). Five-Year Outcomes with Dabrafenib plus Trametinib in Metastatic Melanoma. N. Engl. J. Med. 381, 626–636.

Rodriguez, J., Ren, G., Day, C.R., Zhao, K., Chow, C.C., and Larson, D.R. (2019). Intrinsic Dynamics of a Human Gene Reveal the Basis of Expression Heterogeneity. Cell 176, 213–226.e18.

Rodriguez-Fraticelli, A.E., Weinreb, C., Wang, S.-W., Migueles, R.P., Jankovic, M., Usart, M., Klein, A.M., Lowell, S., and Camargo, F.D. (2020). Single-cell lineage tracing unveils a role for TCF15 in haematopoiesis. Nature 583, 585–589.

Roesch, A., Fukunaga-Kalabis, M., Schmidt, E.C., Zabierowski, S.E., Brafford, P.A., Vultur, A., Basu, D., Gimotty, P., Vogt, T., and Herlyn, M. (2010). A temporarily distinct subpopulation of slow-cycling melanoma cells is required for continuous tumor growth. Cell 141, 583–594.

Roesch, A., Vultur, A., Bogeski, I., Wang, H., Zimmermann, K.M., Speicher, D., Körbel, C., Laschke, M.W., Gimotty, P.A., Philipp, S.E., et al. (2013). Overcoming intrinsic multidrug resistance in melanoma by blocking the mitochondrial respiratory chain of slow-cycling JARID1B(high) cells. Cancer Cell 23, 811–825.

Rutherford, S.L., and Lindquist, S. (1998). Hsp90 as a capacitor for morphological evolution. Nature 396, 336–342.

Schuh, L., Saint-Antoine, M., Sanford, E.M., Emert, B.L., Singh, A., Marr, C., Raj, A., and Goyal, Y. (2020). Gene Networks with Transcriptional Bursting Recapitulate Rare Transient Coordinated High Expression States in Cancer. Cell Syst 10, 363–378.e12.

Shaffer, S.M., Dunagin, M.C., Torborg, S.R., Torre, E.A., Emert, B., Krepler, C., Beqiri, M., Sproesser, K., Brafford, P.A., Xiao, M., et al. (2017). Rare cell variability and drug-induced reprogramming as a mode of cancer drug resistance. Nature 546, 431–435.

Shaffer, S.M., Emert, B.L., Reyes Hueros, R.A., Cote, C., Harmange, G., Schaff, D.L., Sizemore, A.E., Gupte, R., Torre, E., Singh, A., et al. (2020). Memory Sequencing Reveals Heritable Single-Cell Gene Expression Programs Associated with Distinct Cellular Behaviors. Cell 182, 947–959.e17.

Shakiba, N., Fahmy, A., Jayakumaran, G., McGibbon, S., David, L., Trcka, D., Elbaz, J., Puri, M.C., Nagy, A., van der Kooy, D., et al. (2019). Cell competition during reprogramming gives rise to dominant clones. Science 364.

Sharma, S.V., Lee, D.Y., Li, B., Quinlan, M.P., Takahashi, F., Maheswaran, S., McDermott, U., Azizian, N., Zou, L., Fischbach, M.A., et al. (2010). A chromatin-mediated reversible drug-tolerant state in cancer cell subpopulations. Cell 141, 69–80.

Spencer, S.L., Gaudet, S., Albeck, J.G., Burke, J.M., and Sorger, P.K. (2009). Non-genetic origins of cell-to-cell variability in TRAIL-induced apoptosis. Nature 459, 428–432.

Su, Y., Wei, W., Robert, L., Xue, M., Tsoi, J., Garcia-Diaz, A., Homet Moreno, B., Kim, J., Ng, R.H., Lee, J.W., et al. (2017). Single-cell analysis resolves the cell state transition and signaling dynamics associated with melanoma drug-induced resistance. Proc. Natl. Acad. Sci. U. S. A. 114, 13679–13684.

Symmons, O., and Raj, A. (2016). What’s Luck Got to Do with It: Single Cells, Multiple Fates, and Biological Nondeterminism. Mol. Cell 62, 788–802.

Symmons, O., Chang, M., Mellis, I.A., Kalish, J.M., Park, J., Suszták, K., Bartolomei, M.S., and Raj, A. (2019). Allele-specific RNA imaging shows that allelic imbalances can arise in tissues through transcriptional bursting. PLoS Genet. 15, e1007874.

Tian, L., Tomei, S., Schreuder, J., Weber, T.S., Amann-Zalcenstein, D., Lin, D.S., Tran, J., Audiger, C., Chu, M., Jarratt, A., et al. (2021). Clonal multi-omics reveals Bcor as a negative regulator of emergency dendritic cell development. Immunity 54, 1338–1351.e9.

Tirosh, I., Izar, B., Prakadan, S.M., Wadsworth, M.H., 2nd, Treacy, D., Trombetta, J.J., Rotem, A., Rodman, C., Lian, C., Murphy, G., et al. (2016). Dissecting the multicellular ecosystem of metastatic melanoma by single-cell RNA-seq. Science 352, 189–196.

Torre, E.A., Arai, E., Bayatpour, S., Jiang, C.L., Beck, L.E., Emert, B.L., Shaffer, S.M., Mellis, I.A., Fane, M.E., Alicea, G.M., et al. (2021). Genetic screening for single-cell variability modulators driving therapy resistance. Nat. Genet. 53, 76–85.

Umkehrer, C., Holstein, F., Formenti, L., Jude, J., Froussios, K., Neumann, T., Cronin, S.M., Haas, L., Lipp, J.J., Burkard, T.R., et al. (2021). Isolating live cell clones from barcoded populations using CRISPRa-inducible reporters. Nat. Biotechnol. 39, 174–178.

Waddington, C.H. (2014). The Strategy of the Genes (Routledge).

Weinreb, C., Rodriguez-Fraticelli, A., Camargo, F., and Klein, A.M. (2018). Lineage tracing on transcriptional landscapes links state to fate during differentiation.

Wouters, J., Kalender-Atak, Z., Minnoye, L., Spanier, K.I., De Waegeneer, M., Bravo González-Blas, C., Mauduit, D., Davie, K., Hulselmans, G., Najem, A., et al. (2020). Robust gene expression programs underlie recurrent cell states and phenotype switching in melanoma. Nat. Cell Biol. 22, 986–998.

Yunusova, A.M., Fishman, V.S., Vasiliev, G.V., and Battulin, N.R. (2017). Deterministic versus stochastic model of reprogramming: new evidence from cellular barcoding technique. Open Biol. 7.

Zorita, E., Cuscó, P., and Filion, G.J. (2015). Starcode: sequence clustering based on all-pairs search. Bioinformatics 31, 1913–1919.

