## Supplementary Figures for "Pre-determined diversity in resistant fates emerges from homogenous cells after anti-cancer drug treatment"

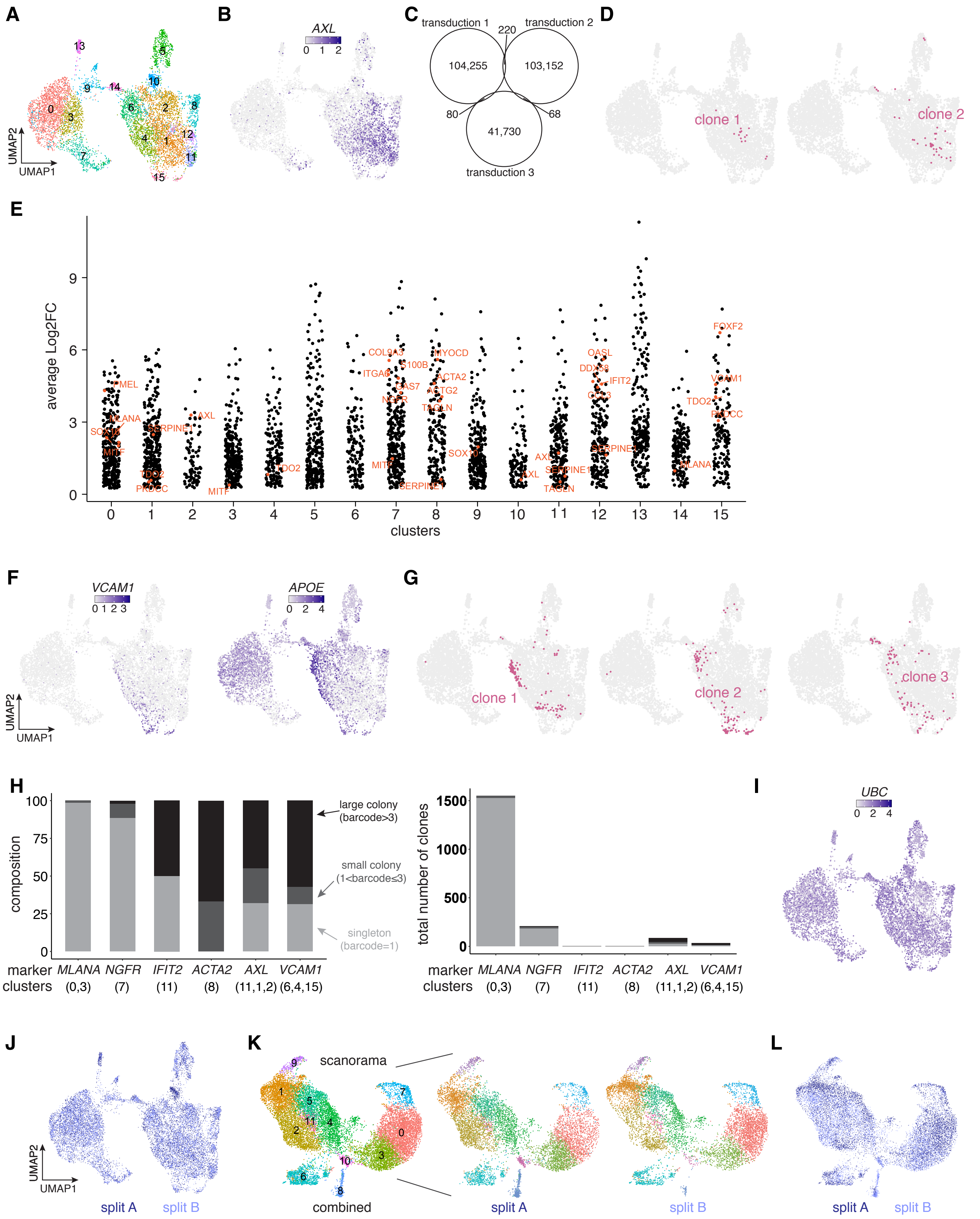


**Supplementary Figure 1**

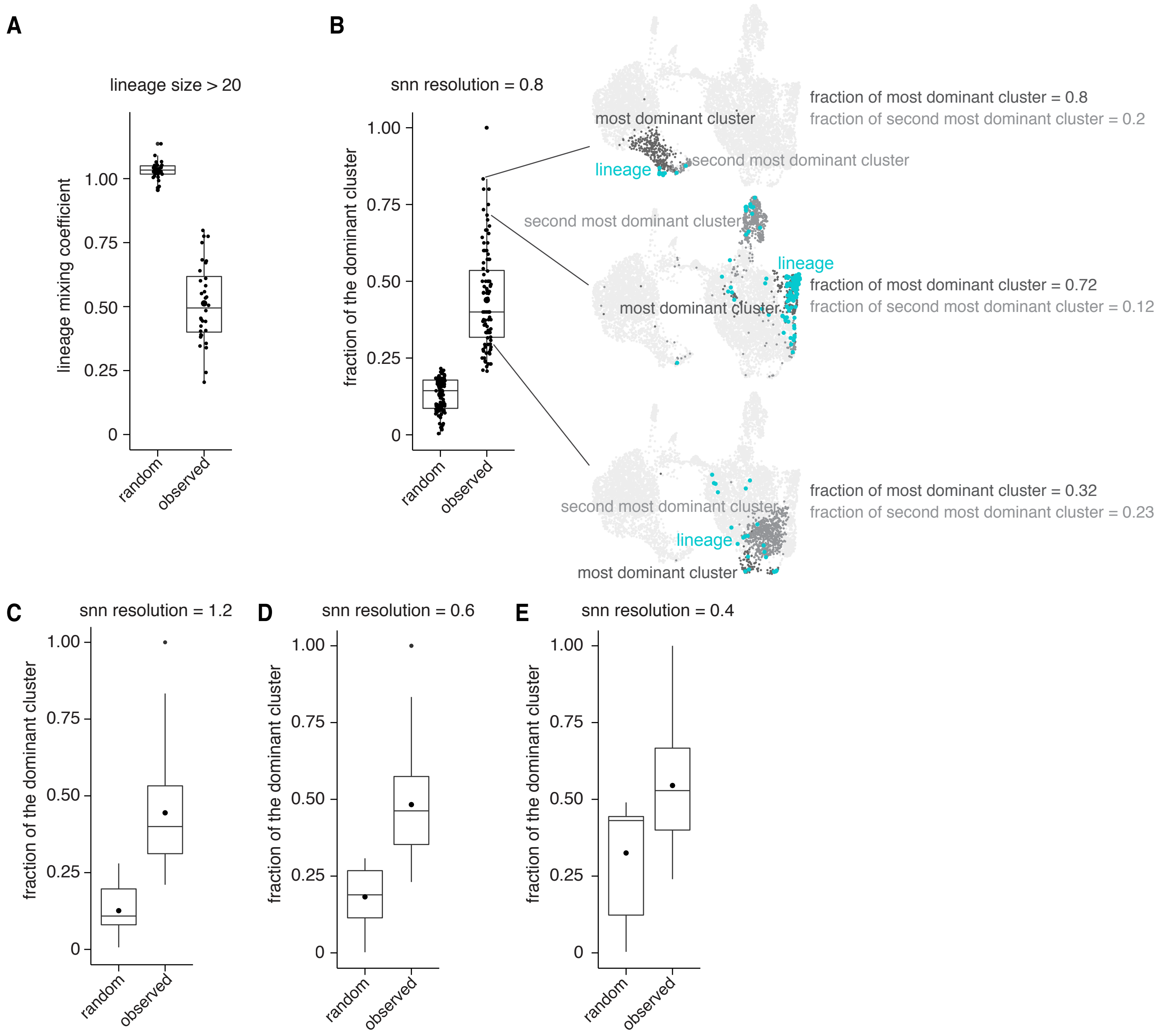


**Supplementary Figure 2**

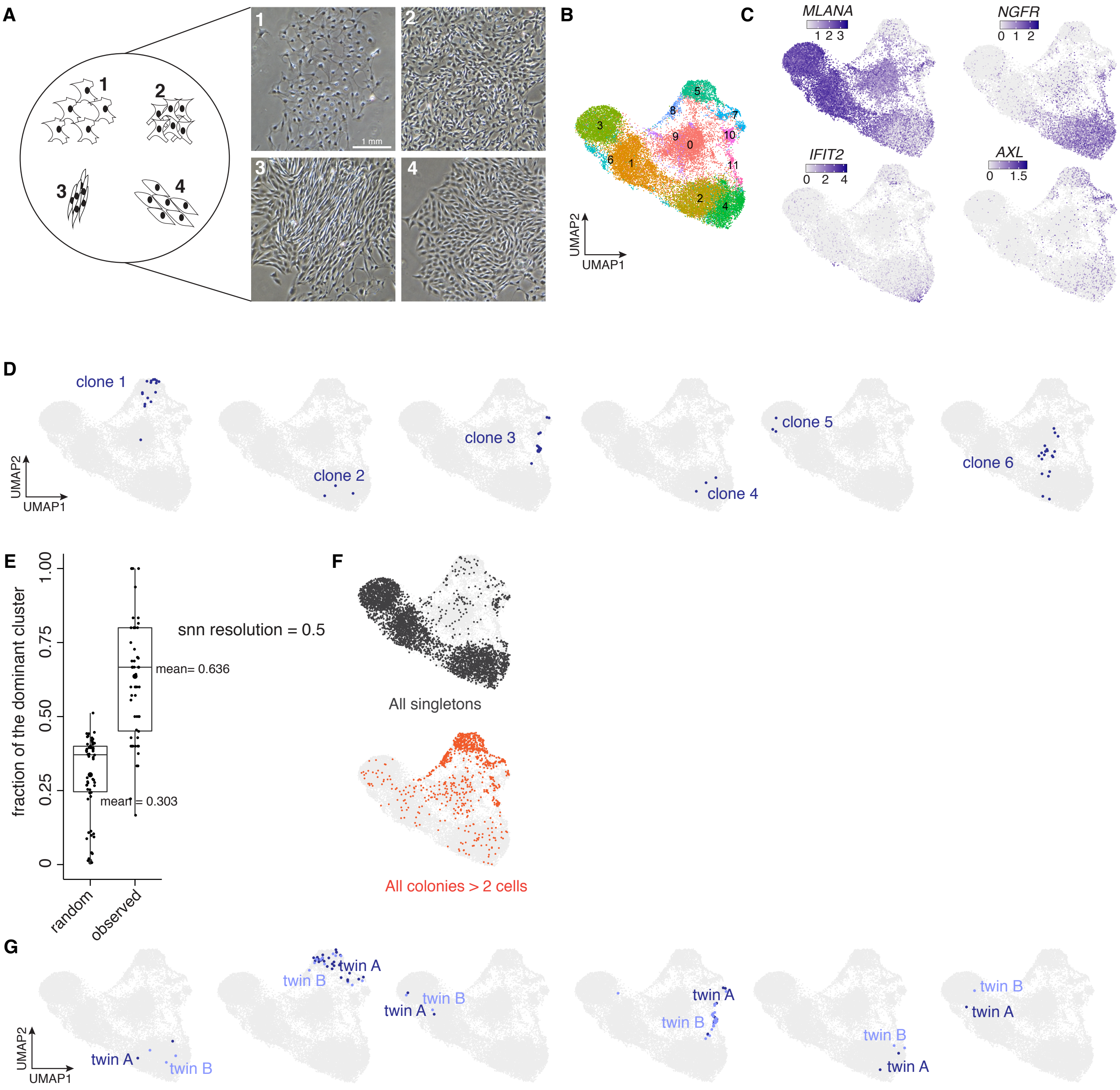


**Supplementary Figure 3**

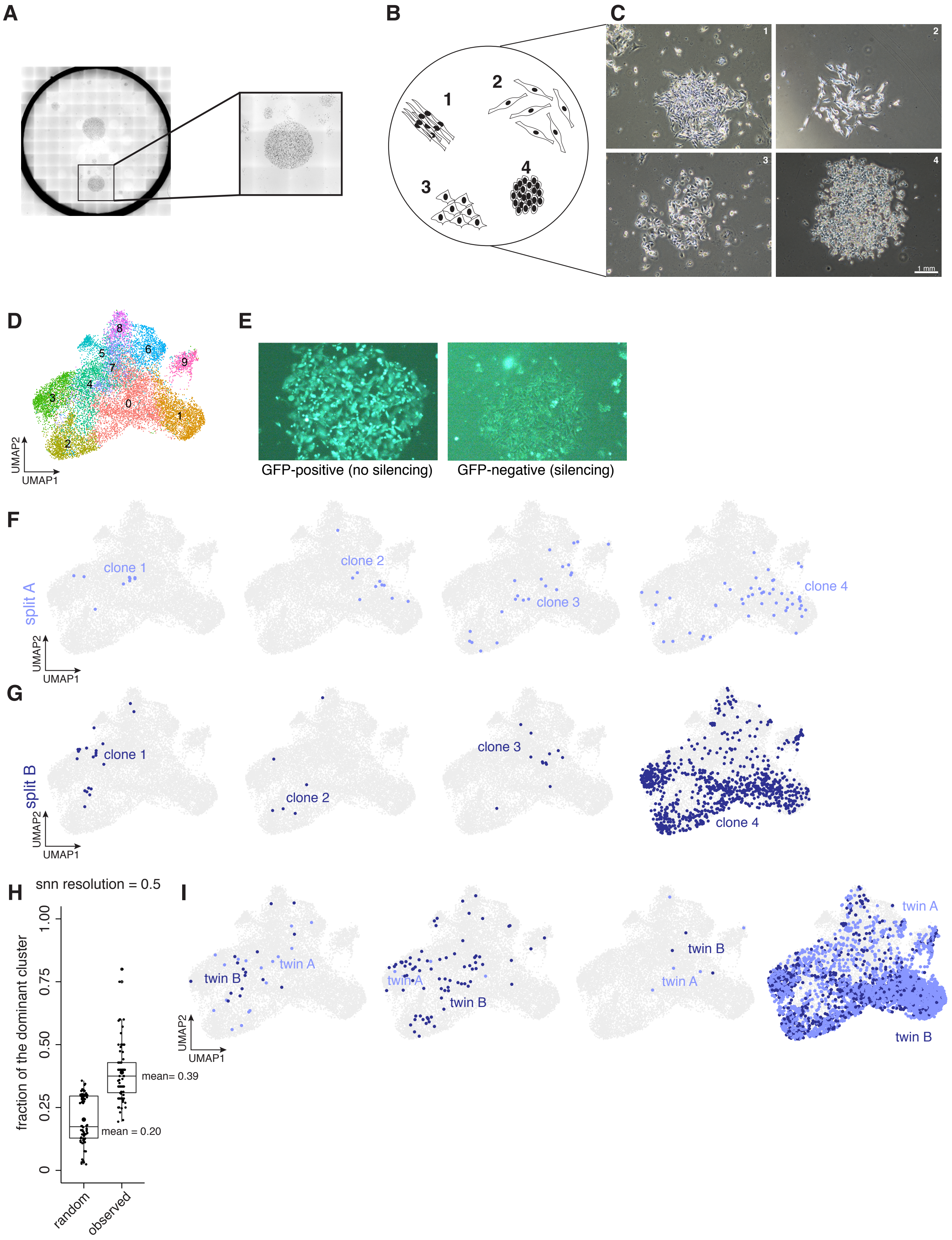


**Supplementary Figure 4**

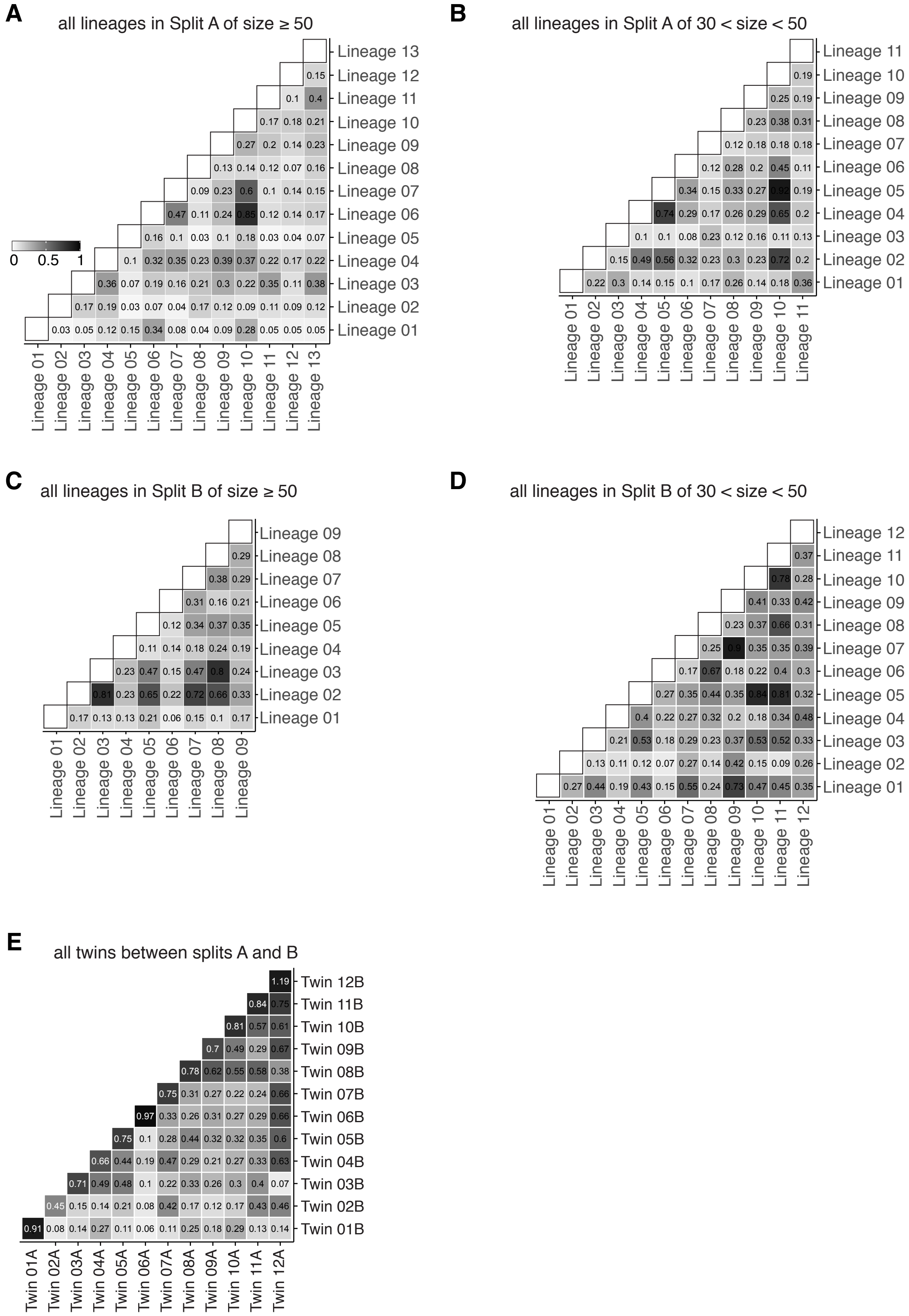


**Supplementary Figure 5**

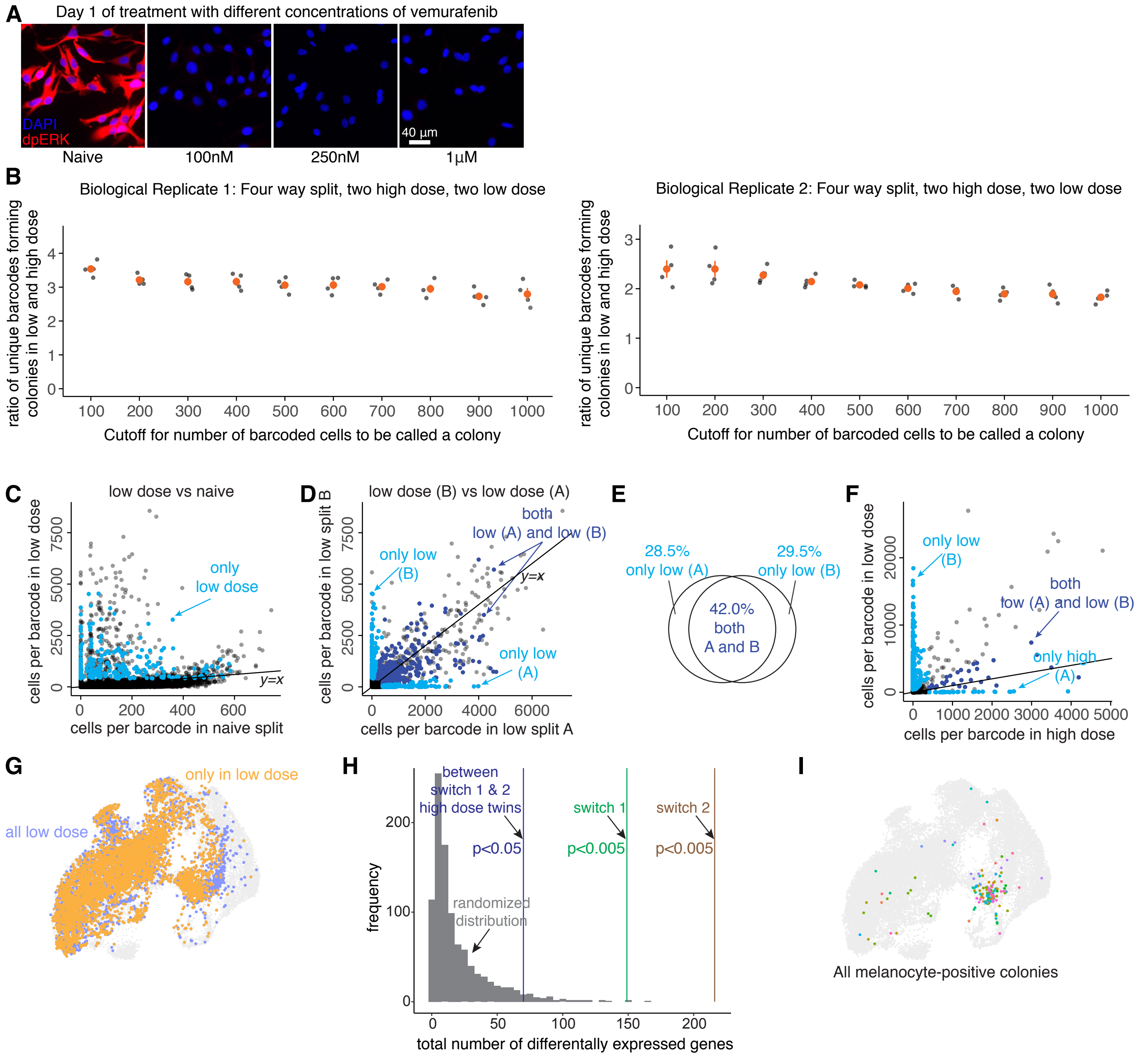


**Supplementary Figure 6**

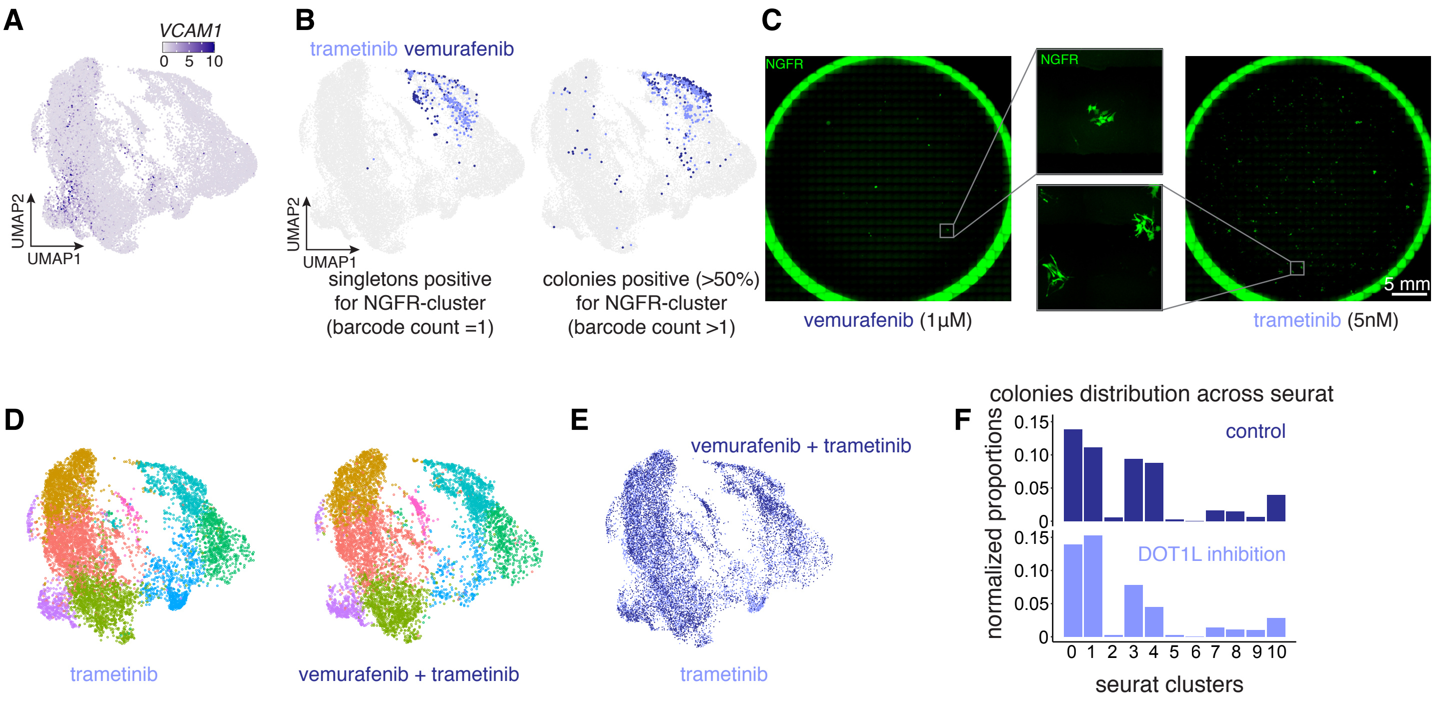


**Supplementary Figure 7**

### Supplementary Figure Captions

**Supplementary Figure 1:** related to Figures 1,2,3, and 4

a. We applied the Uniform Manifold Approximation and Projection (UMAP) algorithm within Seurat to the first 50 principal components to visualize differences in gene expression. Cells are colored by clusters determined using Seurat's FindClusters command at a resolution of 0.6 (i.e. "Seurat clusters, resolution = 0.6").
b. On the UMAP, we recolored each cell by its expression for a select subset of genes that were identified as differentially expressed via the Seurat pipeline and marked different clusters. *AXL*, which is a canonical resistance marker, was found spread across Seurat clusters 1,2, and 11 (snn 0.6).
c. A Venn diagram measuring overlap of barcodes between three independent transductions. No barcode overlap was observed across all three transductions.
d. Two examples demonstrate that clones high for the canonical resistance marker *AXL (*and *SERPINE1)* are constrained largely in specific clusters.
e. Differentially expressed genes per cluster were determined using Seurat's FindAllMarkers command at a resolution of 0.6.
f. *VCAM1* and *APOE* are predominantly present in discontiguous clusters 15 and 6 respectively on UMAP.
g. Three clones high for *VCAM1* (cluster 15) are also high for *APOE* (cluster 6).
h. (left) Markers for different clusters exhibited different proliferative capacities. For example clusters 0 and 3, which are enriched for the gene *MLANA,* predominantly (98.6%) contained singletons. Cluster 7, enriched for the gene *NGFR*, was largely composed of singletons or small colonies. Other markers, including *ACTA2*, *AXL*, and *VCAM1*, which belonged to different clusters, primarily contained small and large colonies, and relatively fewer singletons. (right) UMAP where cells are colored by clusters determined using Seurat's FindClusters command at a resolution of 0.6 (i.e. "Seurat clusters, resolution = 0.6"). (right) Absolute number of clones marking resistant fate clusters and their associated proliferative capacities. Because a large fraction of the clones in clusters 0 and 3 exist as singletons, these clusters contain the most number of unique surviving clones.
i. *UBC*, a housekeeping gene, is expressed ubiquitously throughout the UMAP.
j. UMAP recolored for combined resistant cells from splits A (dark blue) and B (light blue) of the same drug condition of vemurafenib. The two splits are interspersed on the UMAP.
k. UMAP obtained from applying another data integration method, scanorama, on identically-treated samples (split A and B) which produced qualitatively similar results, such that the two splits had a similar distribution of cells in each of the single-cell clusters.
l. UMAP obtained from scanorama is recolored for combined resistant cells from splits A (dark blue) and B (light blue) for the same drug condition (vemurafenib). The two splits are interspersed on the UMAP.

**Supplementary Figure 2:** related to Figure 1

a. We formulated a quantitative metric, the mixing coefficient, to calculate the transcriptional relatedness of different clones to each other. For each pair of barcoded clones, we calculated the nearest neighbors for each cell in the 50-dimensional principal component space. We then classified the neighbors as "self" if the neighbors are from the same barcode clone or "non-self" if they belong to another barcode clone. The mixing coefficient is the ratio of the averaged fraction of self neighbors to the averaged fraction of non-self neighbors. The higher the mixing coefficient, the higher the transcriptional relatedness of the barcoded clones analyzed (a value of 1 corresponds to perfect mixing, and a value of 0 corresponds to no mixing). We reasoned that for randomly sampled barcode clones, the mixing coefficient will be close to 1 because two randomly sampled clones will have a high likelihood of each being spread out uniformly throughout the principal component space. In contrast, the mixing coefficient will be lower than 1 for experimentally observed clones if they each mark different clusters and are not randomly distributed. We found that this was indeed the case where the mixing coefficient was close to 1 for randomly sampled pseudo-clones, which was much higher than the mixing coefficient for experimentally observed clones, suggesting that each clone is constrained in the transcriptional space.

b. We quantified the preference for a specific cluster across all barcode clones (clone size>4). Specifically, we calculated the fraction of dominant clusters for each clone and found it to be significantly higher than that of randomly selected cells (resolution = 0.8). Representative examples of clones covering different fractions of most and second most dominant clusters.
c. We quantified the preference for a specific cluster across all barcode clones (clone size>4). Specifically, we calculated the fraction of dominant clusters for each clone and found it to be significantly higher than that of randomly selected cells (resolution = 1.2).
d. We quantified the preference for a specific cluster across all barcode clones (clone size>4). Specifically, we calculated the fraction of dominant clusters for each clone and found it to be significantly higher than that of randomly selected cells (resolution = 0.6).
e. We quantified the preference for a specific cluster across all barcode clones (clone size>4). Specifically, we calculated the fraction of dominant clusters for each clone and found it to be significantly higher than that of randomly selected cells (resolution = 0.4).

**Supplementary Figure 3:** related to Figure 1,2, and 4

a. (left) For another single-cell derived melanoma cell line WM983B E9-C6, we traced representative resistant cells in Adobe Illustrator and created cartoon schematics based on visual inspection of orientation and density. (right) Brightfield images of resistant colonies exhibiting different types of morphologies.
b. We applied the Uniform Manifold Approximation and Projection (UMAP) algorithm within Seurat to the first 50 principal components to visualize differences in gene expression. Cells are colored by clusters determined using Seurat's FindClusters command at a resolution of 0.5 (i.e. "Seurat clusters, resolution = 0.5").
c. On the UMAP, we recolored each cell by its expression for a select subset of genes that were identified as differentially expressed via the Seurat pipeline and marked different clusters. *MLANA*, which marks melanocytes, is found largely in clusters 1,3, and 5; *IFIT2*, which marks type-1 interferon signaling, is found largely in cluster 4; *NGFR*, which marks neural crest cells, is found largely in cluster 2 and 4; *AXL*, which is a canonical resistance marker, is found largely in cluster 5 and 7.
d. Six examples to demonstrate that a clone (cells sharing the same barcode) is constrained largely in a specific transcriptional cluster such that cells within a clone are more transcriptionally similar to each other than cells in other clones. Some clones are larger in size than others, and some exist as singletons, meaning they survive vemurafenib treatment but do not necessarily divide while exposed to the drug.

e. We quantified the preference for a specific cluster across all barcode clones (clone size>4). Specifically, we calculated the fraction of dominant clusters for each clone and found it to be significantly higher than that for randomly selected cells. The analysis plotted here is for a cluster resolution of 0.5.
f. Painting of singletons and colonies onto the UMAP demonstrated that singletons and colonies belonged to distinct regions and clusters.
g. UMAPs of representative twin clones (sharing the same barcode) across the two splits A and B. The twins largely end up with the same transcriptional fate, invariant of the clone size. This observation suggests that cells are predestined for distinct resistant fates upon exposure to vemurafenib.

**Supplementary Figure 4:** related to Figure 1,2, and 4

a. Nuclei scans (DAPI-stained) of resistant colonies emerging from treatment of the single-cell derived triple negative breast cancer cell line MDA-MB-231-D4 with 1nM paclitaxel.
b. For the MDA-MB-231-D4 cell line, we traced representative resistant cells in Adobe Illustrator and created cartoon schematics based on visual inspection of orientation and density.
c. Brightfield images of resistant colonies exhibiting different types of morphologies.
d. We applied the Uniform Manifold Approximation and Projection (UMAP) algorithm within Seurat to the first 50 principal components to visualize differences in gene expression. Cells are colored by clusters determined using Seurat's FindClusters command at a resolution of 0.5 (i.e. "Seurat clusters, resolution = 0.5").
e. We observed silencing of the transcribed barcodes in a subset of colonies, as revealed by epifluorescence imaging of the GFP signal. The colony on the left is strongly expressing a GFP signal while the colony on the right has a very dim GFP signal.
f. Four examples from split A demonstrate that a clone (cells sharing the same barcode) is constrained largely in specific UMAP regions such that cells within a clone are more transcriptionally similar to each other than cells in other clones.
g. Four examples from split B demonstrate that a clone (cells sharing the same barcode) is constrained largely in specific UMAP regions such that cells within a clone are more transcriptionally similar to each other than cells in other clones.
h. We quantified the preference for a specific cluster across all barcode clones (clone size>4). Specifically, we calculated the fraction of dominant clusters for each clone and found it to be significantly higher than that for randomly selected cells. The analysis plotted here is for a cluster resolution of 0.5.
i. UMAPs of representative twin clones (sharing the same barcode) across the two splits A and B. The twins largely end up with the same transcriptional fate, invariant of the clone size. This observation suggests that cells are predestined for distinct resistant fates upon exposure to chemotherapy drug paclitaxel.

**Supplementary Figure 5:** related to Figure 4

a. We formulated a quantitative metric, the mixing coefficient, to calculate the transcriptional relatedness of different clones to each other. For each pair of barcoded clones, we calculated the nearest neighbors for each cell in the 50-dimensional principal component space. We then classified the neighbors as "self" if the neighbors are from the same barcode clone or "non-self" if they belong to another barcode clone. The mixing coefficient is the ratio of averaged fraction of self neighbors to averaged fraction non-self neighbors. The higher the mixing coefficient, the higher the transcriptional relatedness of the barcoded clones analyzed (a value of 1 corresponds to perfect mixing, and a value of 0 corresponds to no mixing). Pairwise calculation of the mixing coefficient for all split A clones of size greater than or equal to 50.

b. Pairwise calculation of the mixing coefficient for all split A clones of size between 30 and 50.
c. Pairwise calculation of the mixing coefficient for all split B clones of size greater than or equal to 50.
d. Pairwise calculation of the mixing coefficient for all split B clones of size between 30 and 50.
e. The mixing coefficient for twin and non-twin clones across splits A and B is presented with representative examples on the UMAP.

**Supplementary Figure 6:** related to Figure 5

a. We performed antibody stainings for the active, dually phosphorylated form of ERK on WM989 A6-G3 cells, either naive or treated with different doses of vemurafenib. Treated cells were fixed and imaged after 1 day of treatment. All doses of vemurafenib tested (100nM-1uM) eliminated the Ras/ERK signaling compared to the naive cells.
b. Quantification of the total number of unique barcodes forming colonies from each dose across biological replicates. Since this analysis can be dependent on where we define the cutoff for a barcode to be considered as a colony forming barcode, we used a range of cutoffs to show that the fold-change in unique barcodes for the low dose compared to the high dose treatment remains largely invariant of the cutoff. Therefore, changing the cutoff for minimum cell count does not affect our conclusions.
c. We observed no correlation between the cell count (as measured by gDNA sequencing of the barcodes) of low dose only resistant clones (light blue) and their counterparts in the initial population from another split. This result rules out the possibility that the additional new clones could have been a result of faster dividing cells in the initial population.
d. We compared the clones that only survived at the low dose across two different low dose splits. For this, we collected the barcodes that only survived in two different low dose splits compared to the high dose splits and plotted them against each other. We found that many clones present in only the low dose also tended to show up across the two low dose splits (dark blue), showing that these low-dose only clones are also the result of intrinsic pre-existing differences between initial cells.
e. A venn diagram showing that, amongst the barcodes surviving only in low doses, there is a significant fraction of overlap when comparing across the two low dose splits.
f. gDNA barcode sequencing on the cells remaining after performing FateMap on a subset of resistant cells. For each barcode identified by sequencing, we plotted its abundance in corresponding splits A (high dose) and B (low dose). Those present in both high and low dose splits are colored in dark blue, and those present only in either A (high) or B (low) are colored in cyan. Those present in both (dark blue) exhibited a strong correlation, suggesting that their ability to survive and become resistant is invariant of drug dose. For those present only in either (cyan), we found them to be much more abundant in low dose (split B), suggesting that new barcodes that are otherwise unable to survive in the high dose become drug resistant in the low dose. The high number of cells on each axis compared to on the y=x axis is likely because of losses incurred during the preparation of the FateMap experiment.

g. Painting of all barcode clones on UMAP that were present only in the low dose split B from panel F (yellow), compared to all the low dose resistant clones detected by FateMap. We find that while many of the low dose only clones occupy regions interspersed with high dose cells, there are specific non-overlapping regions as well.
h. Comparison of the total number of differentially expressed genes for three pairwise comparisons: 149 genes for switch 1, 216 genes for switch 2, and 70 genes for comparison within the high dose for switch 1 and switch 2. The total number of differentially expressed genes for each comparison was significantly more than the number of genes for randomly sampled cells from the NGFR-high cluster 9 (simulation ran 1000 times).
i. Painting of colony forming barcodes within the MLANA-high cluster 5, which is largely occupied by cells from the low dose of vemurafenib. Therefore, there are several large colony-forming clones that come from MLANA-high cells, which is not the case for MLANA-high cells in the high dose where these cells largely exist as singletons.

We next wondered if the clones that only survived at low dose were also predetermined to survive low dose. That is, if we exposed two separated twin splits to the same low dose, would the clones that were detected only in low dose show a strong correspondence across low dose splits? We found that the clones present in only the low dose also tended to show up across doses, showing that these low dose-only clones are also the result of intrinsic, pre-existing differences between initial cells as opposed to microenvironmental differences (Supplementary Figure 6C-E).

**Supplementary Figure 7:** related to Figure 6 and 7

a. UMAP for combined vemurafenib and trametinib treatment conditions recolored for each cell by its expression of the gene *MLANA,* which is enriched in cluster 6.
b. Painting of singletons and colonies onto the UMAP for the NGFR-high cluster 4, colored by the condition, showing a relative enrichment of cells from trametinib as compared to vemurafenib. This panel also demonstrates that both singletons and colonies occupy cluster 4 from each of the two conditions.
c. We performed antibody stainings for NGFR on colonies emerging from treatment of the same number of starting melanoma cells with either vemurafenib or trametinib. Consistent with FateMap, we found an increased number of NGFR-positive resistant cells in trametinib treated cells as compared to the vemurafenib treatment.
d. UMAP split by each drug condition (trametinib or vemurafenib and trametinib), with colors representing clusters determined using Seurat's FindClusters command at a resolution of 0.5 (i.e. "Seurat clusters, resolution = 0.5").
e. UMAP recolored for combined resistant cells from trametinib (light blue) and vemurafenib and trametinib (dark blue). The cells from two conditions are interspersed into each other on the UMAP.
f. Distribution of cells across clusters for control (top) and DOT1L inhibitor-pretreated (bottom) conditions for clone size>2.

**Supplementary Movie 1:** Time lapse imaging of single-cell derived WM989 A6-G3 cells exposed to a 1μM dose of vemurafenib. The time stamp is provided in top left, and time 0 represents the time at which the drug was added.

**Supplementary Movie 2:** Time lapse imaging of single-cell derived WM989 A6-G3 cells exposed to a 100nM dose of vemurafenib. The time stamp is provided in top left, and time 0 represents the time at which the drug was added.

**Supplementary Movie 3:** Time lapse imaging of single-cell derived WM989 A6-G3 cells exposed to a 5nM dose of trametinib. The time stamp is provided in top left, and time 0 represents the time at which the drug was added.

**Supplementary Tables:** Captions mentioned within each table tab.
